# Structural basis for size-selective perception of chitin in plants

**DOI:** 10.1101/2025.07.09.663820

**Authors:** Kira Gysel, Simon Boje Hansen, Henriette Rübsam, Husam M.A.B. Alsarraf, Eva Madland, Jeryl Xin Jie Cheng, Caroline Baadegaard, Emil Christian Poulsen, Maria Vinther, Sebastien Fort, Jens Stougaard, Kasper Røjkjær Andersen

## Abstract

Plants detect microbes through pattern recognition receptors that perceive conserved microbial surface motifs known as microbe-associated molecular patterns (MAMPs). LysM receptors recognize and mediate downstream responses to chitinous MAMPs. Here, we elucidate the mechanism for the specific recognition of long-chain chitin oligomers and identify a hallmark bridge domain characteristic for the CHIP receptor class. Structural analysis of receptor-ligand complexes, biochemistry, and *in planta* functional studies using inhibitory nanobodies reveal the mechanism of size-selective, high affinity chitin perception in *Lotus japonicus* and *Medicago truncatula*. Additionally, we identify CERKs as low-affinity, yet essential co-receptors and propose a mechanistic model for a ligand-induced core signaling heterocomplex. Our findings provide mechanistic insights into plant chitin perception and the formation of receptor complexes critical for immune signaling.

## Introduction

The plant immune system relies on pattern recognition receptors to identify microbe-associated molecular patterns (MAMPs) and mount appropriate immune responses towards microbes. The MAMP chitin, a β-1,4 polymer of N-acetylglucosamine, is a major constituent of the fungal cell wall and is detected by the extracellular lysin motifs (LysM) of LysM receptor kinases (LysM-RK) and LysM receptor proteins (LysM-RP) (*1*, *2*).

LysM-RKs with an active intracellular kinase domain induce chitin-triggered signaling in complex with LysM receptors that either have a pseudokinase or lack the kinase domain entirely (LysM-RPs). Earlier studies have demonstrated that longer chitin oligomers, especially octa-N-acetyl-chitooctaose (CO8), have increased immunogenicity compared to shorter chitin oligomers, such as penta-N-acetyl-chitopentaose (CO5) (*3–6*). Together with fungal lipochitooligosaccharides, chitin fragments, especially short chain, are also important signaling molecules for the establishment of arbuscular mycorrhizal symbiosis (AMS). Stringent distinction between chitinous MAMPs from pathogenic and symbiotic origins is critical for many plants, although the molecular mechanisms behind this are not fully understood (*4*, *6*, *7*).

In *Arabidopsis thaliana (Arabidopsis),* the LysM-RK *At*CERK1 contains an active kinase and is essential for all chitin-elicited defense responses (*8*). *At*CERK1 functions in complex with the LysM-RK pseudokinases *At*LYK4 and *At*LYK5 that are partially redundant, as only *lyk4 lyk5* double knockout plants are unable to mount chitin-elicited immunity (*9–11*). *At*CERK1, as well as *At*LYK4 and *At*LYK5, are high-affinity chitin binders, while in rice and *Marchantia paleacea* the CERK-type receptors have low chitin affinity compared to their LysM-RK pseudokinase or LysM-RP partner (*11–16*).

In the model legumes *Lotus japonicus* (*Lotus*) and *Medicago truncatula (Medicago)*, the active kinases *Lj*CERK6 (from here on CERK6) and *Mt*CERK1 as well as the pseudokinases *Lj*LYS13/*Lj*LYS14 (a tandem gene duplication with >90% identity, here renamed to CHIP13/CHIP14 reflecting their function in *Ch*itin *P*erception) and *Mt*LYR4 (from here on LYR4) were identified as pattern recognition receptors involved in chitin-elicited defense (*3–5*, *17*). While the ligand binding mechanisms of CERK-type receptors was studied previously, no structural data is available for CHIP-type LysM-RKs (LYR-III class, (*18*) to understand their role and functions in plant immunity.

Here, we elucidate the structural basis for size selective and high-affinity perception of long chitin oligomers by CHIP-type LysM-RKs. In contrast, legume CERK receptors show comparatively lower chitin affinity but are nevertheless essential in chitin-elicited defense. Using receptor mutants and inhibitory nanobodies we demonstrate how CHIP- and CERK-type LysM-RKs perceive chitin oligomers, facilitating the formation of a signaling-competent receptor complex. Based on our findings, we propose a model in which CHIP13 receptor dimers bind long chain chitin oligomers, enabling ligand-mediated recruitment of CERK6 receptors to initiate immune signaling.

### CHIP13 is a high-affinity receptor for long chitin oligomers

To investigate the molecular mechanisms of chitin perception and especially the role of the pseudokinase receptors CHIP13/14, we recombinantly expressed the extracellular domain of CHIP13 (residues 33-253) in insect cells (Figure S1) and assayed binding towards chitin oligomers with different degrees of polymerization. Quantitative equilibrium dissociation constant (K_d_) measurements using isothermal titration calorimetry (ITC) revealed that CHIP13 binds CO8 with K_d_= 0.9 µM, but CO5 only weakly with K_d_=1.27 mM, a difference of more than 1000-fold (Fig. 1A, fig. S1D-G).

**Fig. 1.**
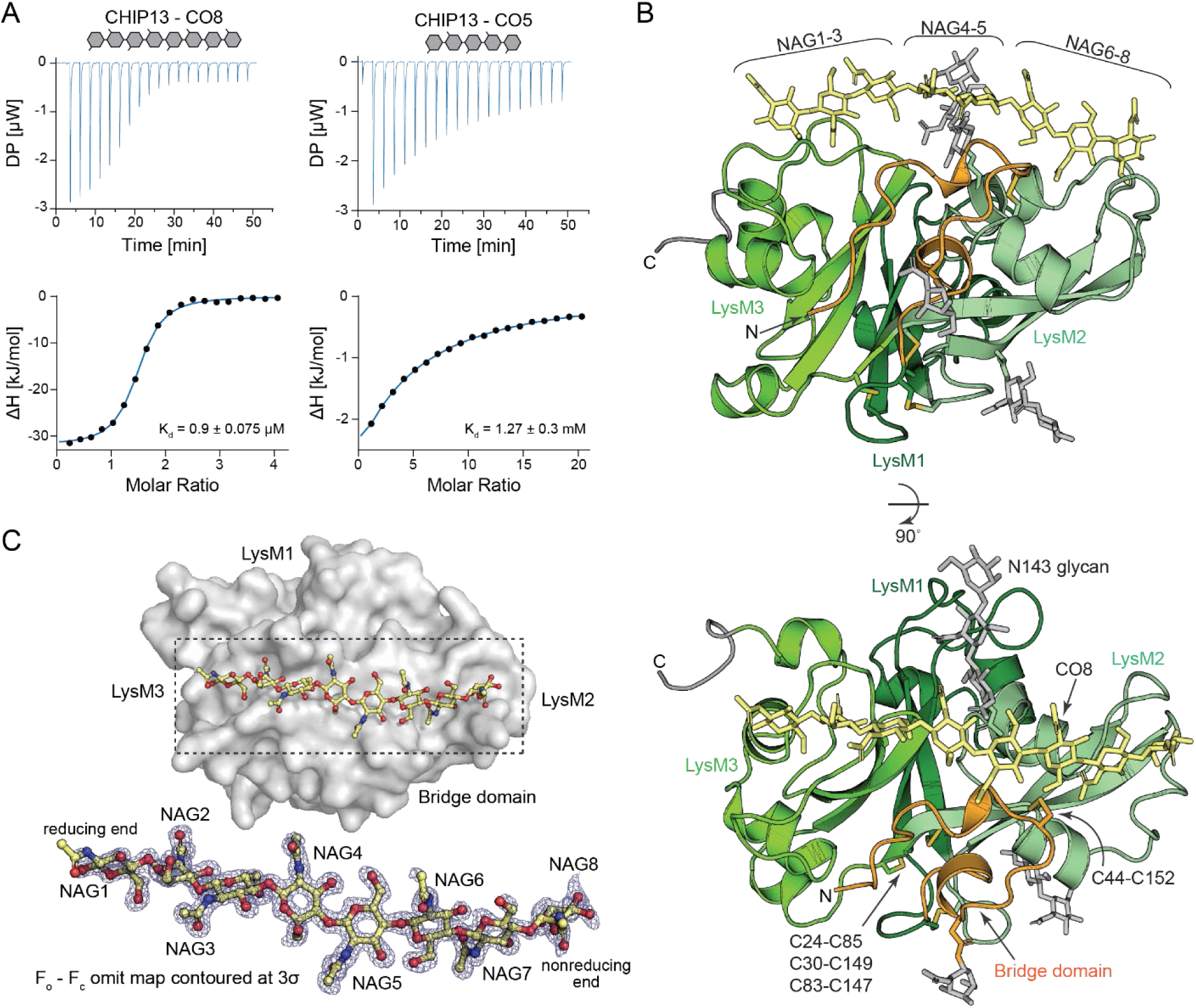
CHIP13 is a size-selective high affinity receptor for long chitin oligomers. (A) ITC isotherms for titration of CO8 (left) and CO5 (right) into CHIP13. (B) Cartoon representation of the crystal structure of the CHIP13:CO8 complex displayed from two angles. The bridge domain is orange, LysM1-3 in different shades of green. The CO8 ligand, counted from the reducing end, is highlighted in yellow stick representation, N-glycans as gray sticks. Conserved LysM-RK disulphide bridges and the additional C44-C152 disulphide are displayed as yellow sticks. (C) F_o_ – F_c_ omit map of CO8 electron density contoured at 3σ. Resolution is d =1.35 Å. Top panel shows CO8 as ball-and-sticks bound by CHIP13 in surface representation. Dashed box indicates ligand and ligand binding site.

To elucidate the molecular mechanisms for this size selective binding of long chitin oligomers, we determined the crystal structure of the CHIP13 ectodomain in complex with CO8 (Fig. 1B, fig. S3C-E, table S1). Analogous to other LysM receptors, CHIP13 forms a globular structure with three canonical LysM domains in a cloverleaf-like arrangement around a central β-sheet connected by a conserved triple set of disulfide bridges (*3*, *5*, *14*, *16*). Unexpectedly, the N-terminal part of CHIP13 (residues 33-55) forms a distinct domain, which is anchored to the LysM2 α2 through an additional disulfide bond (C44-C152) (fig. S1H, I). CO8 is well-defined in the electron density (Fig. 1C) and forms contacts with LysM2, LysM3, and the N-terminal domain. Starting from the reducing end, N-acetylglucosamines 1-3 (NAG1-3) are bound by LysM3 and NAG 6-8, at the nonreducing end, are bound by LysM2. The central part of CO8, NAG4-5, is contacted by the proximal core NAG of the N143 N-glycan and the distinct N-terminal domain, which ‘bridges’ CO8 between LysM2 and LysM3. Hence, we refer to it as the “bridge domain”. This novel domain can be structurally predicted in many LysM-RKs in higher plants and is one of the hallmark features of CHIP chitin receptors (fig. S2).

The collaborative CO8 binding by the two LysM and the bridge domain explains size-selectivity, as shorter chitins cannot reach across the bridge domain. CHIP13 has a defined binding mode for CO8, as the distal exit of LysM2 is blocked by F167. This residue prevents chitin ‘sliding’, which is frequently observed in other LysM proteins. In contrast, the distal exit of LysM3 is open, making accommodation of even longer chitin oligomers than CO8 potentially possible (fig. S3A,B).

### Carbohydrate-π interactions give high ligand affinity and signaling functionality

Having defined the CHIP13 binding site, the mechanism for size-selectivity and the ligand binding mode, we investigated whether the high affinity for CO8 is mediated solely by the increased avidity provided by the two LysM and bridge domains. LysM2 and LysM3 bind CO8 with an extended H-bond network, engaging mostly with the protein backbone, in the shallow grooves formed by the loops between β1-α1 and α2-β2. This binding mode is typical for LysM proteins (*16*, *19–21*). Comparison of the ligand-bound and unbound CHIP13 crystal structures shows that the general fold remains unchanged upon ligand binding (fig. S3B,G). However, two aromatic residues in LysM3 at the interface of the bridge domain, Y232 and F233, as well as the proximal NAG of the N-glycan on N143 in the bridge domain change rotamer and are directly involved in CO8 binding (Fig. 2A). While unaligned in the structure without ligand, these residues form a carbohydrate-π (CH-π) interaction network with both the ligand directly and the N143 glycan, which in turn directly contacts CO8 (Fig. 2B). To investigate the contribution of the CH-π network to binding affinity we generated the CHIP13 Y232A F233A variant, which disrupts the CH-π interaction network while maintaining the LysM2 and LysM3 binding sites (fig. S3H). CHIP13 Y232A F233A has a 200-fold reduced affinity to CO8 (Fig. 2C), supporting the major contribution of CH-π interactions to the high chitin binding affinity of CHIP13.

**Fig. 2.**
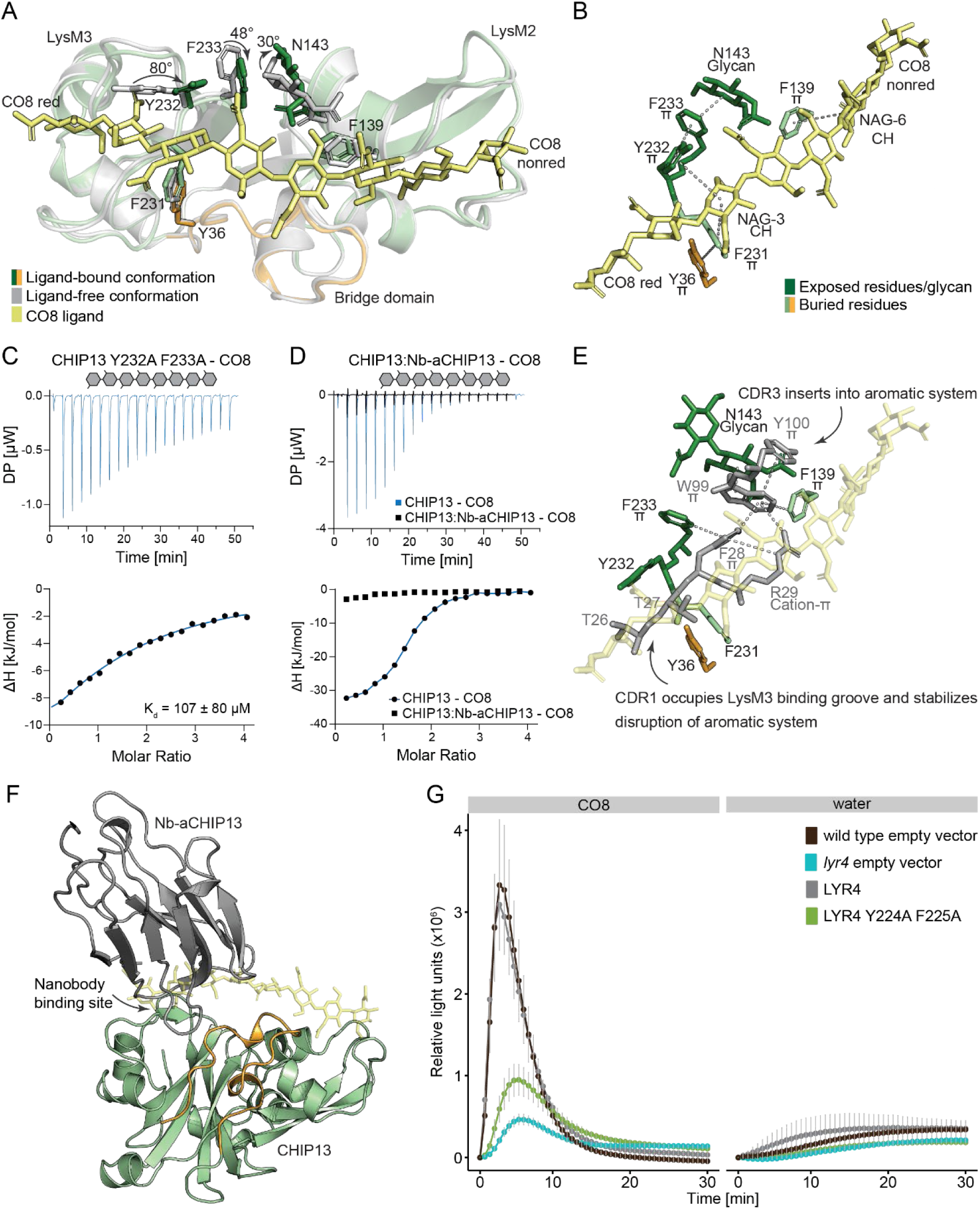
Carbohydrate-π interactions mediate high affinity chitin binding and are essential for ROS production. (A) Superposition of ligand-free CHIP13 (gray) and CHIP13:CO8 complex (light green and orange). For clarity, LysM1 is not displayed. CO8 from CHIP13:CO8 is displayed in yellow stick representation. Residues changing rotamer upon ligand binding are indicated in dark green. Buried or partially buried aromatic residues associated with carbohydrate binding and not changing rotamer are displayed in light green and orange sticks, respectively. (B) Detailed close-up of the CH-π network mediating high binding affinity. Residues engaging in CH-π interactions are displayed in stick representation and indicated with CH (NAGs) or π (aromatic side chains), respectively. Critical interactions are indicated with dashed lines. (C) ITC isotherm for titration of CO8 into CHIP13 Y232A F233A. (D) ITC isotherm for titration of CO8 into the purified CHIP13:Nb-aCHIP13 complex. A replicate of the isotherm for titration of CO8 into CHIP13 is included for comparison. (E) Detailed close-up of the CHIP13:Nb-aCHIP13 binding site. Nb-aCHIP13 is displayed in gray stick representation. Critical interactions for inhibition of chitin binding by Nb-aCHIP13 are shown with dashed lines. (F) Overview of the CHIP13:Nb-aCHIP13 complex with transparent CO8 ligand superimposed from the CHIP13:CO8 crystal structure (G) ROS production in *Medicago* wild type and *lyr4* single mutant roots transformed with *Lyr4* or *Lyr4* Y224A F225A over a period of 30 minutes after treatment with 1 µM CO8 or water (mean ± standard error of the mean, n = 12-16 (CO8) or n = 4-7 (water)).

We raised a high-affinity nanobody against purified CHIP13 (Nb-aCHIP13, K_d_ = 8.1 nM, Figure S4) and observed that Nb-aCHIP13 bound CHIP13 is incapable of chitin binding (Fig. 2D). Also, Nb-aCHIP13 cannot bind CHIP13 Y232A F233A (fig. S4A-C), and our crystal structure of the CHIP13:Nb-aCHIP13 complex confirms that the nanobody directly targets the CO8 binding site of CHIP13. Nb-aCHIP13 blocks LysM3 by occupying its binding groove with complementarity-determining region (CDR) 1 and inserts CDR1 and CDR3 into the CHIP13 aromatic system (Fig. 2E,F, fig. S3D). We conclude that the CO8 binding mode mediated by LysM2, LysM3 and bridge domains *in crystallo*, is the same *in vitro* binding mode assayed by ITC.

Our next aim was to assay the functional relevance of the CH-π network *in planta*. In *Lotus*, the *Chip13/Lys13* and *Chip14*/*Lys14* genes are functionally redundant tandem duplicates in close genomic proximity, and we were unable to generate a double mutant from the LORE1 population (*3*, *22*, *23*). We therefore characterized and conducted mutant studies in *Medicago* with its single ortholog *Lyr4* (*3*, *17*). Sequence analysis and our crystal structure of the LYR4 ectodomain show high similarity to CHIP13. Both the bridge domain, the additional disulfide linking the domain with LysM2, and the aromatic system in LysM3 are present. *In silico* docking of CO8 into the LYR4 crystal structure suggests chitin is bound through the same mechanism as seen in the CHIP13:CO8 crystal structures (fig. S5A-D). ITC assays using purified LYR4 ectodomain reveal a similar high affinity for long chitin oligomers (K_d, CO8_= 1.92 µM) and a 50-fold lower affinity for short chitins (K_d, CO5_= 96 µM) (fig. S5E-F). Previously, we showed that *lyr4* plants exhibit only minor reactive oxygen species (ROS) production when treated with chitin (*3*, *4*, *17*). Complementation of *lyr4* with wild-type *Lyr4* restores ROS production in response to chitin, whereas the *Lyr4 Y224A F225A* variant is significantly impaired (Fig. 2G). Together, this underlines the functional significance of the CH-π network and shows that the high affinity and size-selective chitin binding of LYR4 and CHIP13 is critical for their function in chitin-elicited defense.

### Legume CERK receptors are weak chitin binders and mediate chitin-elicited immunity with CHIP13-type receptors

Following characterization of the CHIP13-type receptors, we investigated the role of the co-receptor CERK6 and its *Medicago* ortholog, *Mt*CERK1 (*3*, *4*, *8*, *11*). Purified CERK6 and *Mt*CERK1 ectodomains show only weak affinity for chitin (2.8 mM and 729 µM for CO5, respectively), to the point that we could not confidently fit the CO8 binding due to limited ligand solubility (Fig. 3A, fig. S6A). In addition, neither CERK6 nor *Mt*CERK1 are retained in a chitin affinity PAGE experiment, whereas CHIP13 and LYR4 show strong retention (fig. S6B). This is in line with reports on *Os*CERK1 that has millimolar affinity for chitin oligomers but is nevertheless essential for chitin-elicited defense in rice (*20*, *24*). In contrast, the *At*CERK1 ectodomain is retained in the chitin affinity gel, aligning with previous studies that it binds chitin with higher affinity than other CERK receptors (fig. S6B) (*9*, *14*, *16*).

**Fig. 3.**
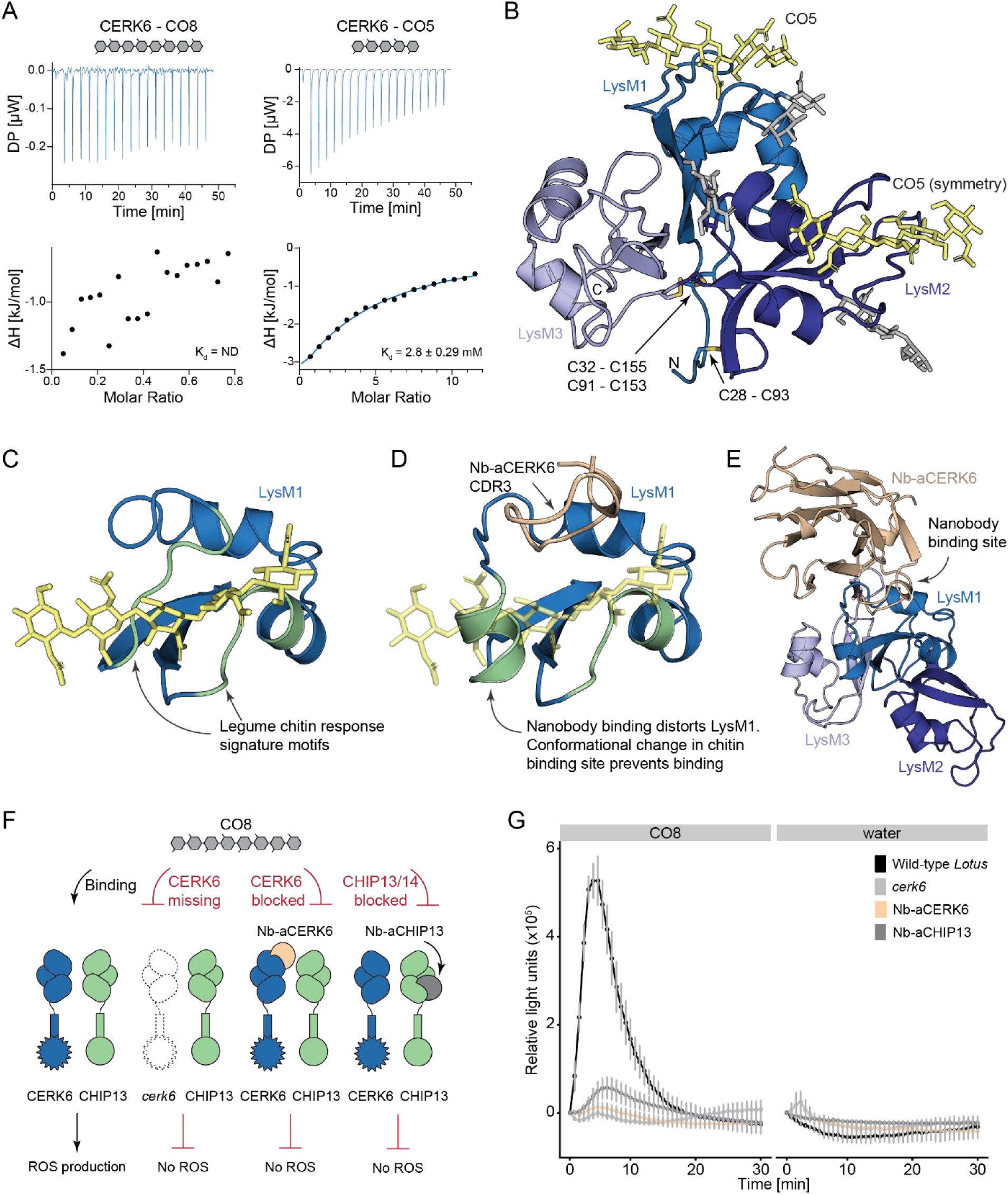
CERK6 is a weak chitin binder and mediates chitin-induced ROS production together with CHIP13. (A) ITC isotherm for titration of CO8 (left) and CO5 (right) into CERK6. Measurement of CO8 binding to CERK6 is limited by CO8 solubility. (B) Cartoon representation of the CERK6 crystal structure in complex with CO5. N-glycans are displayed in gray, the CO5 ligand in yellow stick representation. The ligand is accommodated in LysM1 and LysM2. (C) Close-up of LysM1 and chitin in the CERK6:CO5 crystal structure. Legume chitin response signature motifs in LysM1 are highlighted in green and directly involved in ligand binding. (D) Close-up of LysM1 and CDR3 from the CERK6:Nb-aCERK6:Nb-aCERK6-2 crystal structure. CO5 ligand from CERK6:CO5 structure is superpositioned in semi-transparent stick representation. Nb-aCERK6 blocks the chitin binding site by distorting the LysM1 conformation. (E) Overview of CERK6:Nb-aCERK6 binding mode in cartoon representation. Nb-aCERK6 (beige) binds CERK6 in LysM1. (F) Schematic of nanobody assay on *Lotus* seedlings. (G) ROS production in *Lotus* wild-type seedlings (untreated wild-type, untreated *cerk6*, wild-type pretreated with Nb-aCERK6 or Nb-aCHIP13, respectively) over a period of 30 min after elicitation with 1 µM CO8 (left) or water (mean ± standard error of the mean, n = 5-10 (CO8) or n = 2-4 (water)).

To determine the chitin binding mode of CERK6, we co-crystallized the CERK6 ectodomain with CO5 (Fig. 3B, fig. S6C). The crystal structure shows that CERK6 accommodates chitin in both LysM1 and LysM2 similarly to the previously published crystal structure of *Os*CERK1, including the sandwich-type binding of a single chitin molecule by both LysM1 and a symmetry-related LysM2 (fig. S6E-G, (*20*). Our crystal structure of apo *Mt*CERK1 reveals a conserved fold and both LysM1 and LysM2 can accommodate the CERK6 ligand *in silico* (fig. S6D). Previously, we identified conserved legume chitin response motifs in LysM1 which are essential for the function of legume CERK-type receptors (*5*). Our structures confirm that these signature motifs correspond to the two loops that form the chitin binding site and directly interact with CO5 (Fig. 3C, S6F).

We raised two high affinity nanobodies against the CERK6 ectodomain and isolated a complex of CERK6 bound by both nanobodies simultaneously (fig. S7A, B). From the crystal structure of this complex, we find that one nanobody (Nb-aCERK6) contacts LysM1, distorts the conformation of one of the chitin binding loops and consequently prevents chitin from binding in LysM1 (Fig. 3D). Due to the low chitin binding affinity of CERK6, we were not able to measure the effect of Nb-aCERK6 on chitin binding with ITC. Using saturation transfer difference nuclear magnetic resonance, we show that CERK6 retains some chitin binding capacity in LysM2 when in complex with Nb-aCERK6 (fig. S8), validating that CERK6 in solution is capable of accommodating chitin in both LysM1 and LysM2.

Next, we probed *Lotus* chitin immune signaling *in planta* using our specific nanobodies. Both Nb-aCHIP13 and Nb-aCERK6 directly target the receptors’ chitin binding sites and are strong competitive inhibitors, exceeding *in vitro* chitin binding affinity by orders of magnitude. We pre-treated *Lotus* wildtype seedlings with purified Nb-aCERK6 and observed significant attenuation of chitin-elicited ROS production upon challenge with CO8 comparable to the *cerk6* phenotype (Fig. 3E,F), validating that only LysM1 is required for CERK6s function in immunity (*5*). Similarly, Nb-aCHIP13 treated *Lotus* seedlings showed strong attenuation of chitin-elicited ROS production, supporting that Nb-aCHIP13 targets both CHIP13 and CHIP14. Both receptors share a 93% amino acid identity and the Nb-aCHIP13 epitope is conserved in CHIP14 (Fig. 3E, F, fig. S4D). Our nanobody assays show that CHIP13/14 and CERK6 are essential for chitin-elicited defense in *Lotus* and the target epitopes on the respective receptors are required for ligand perception and immune signaling.

### CHIP13 and CERK6 associate into a receptor complex upon ligand binding

Our results show that chitin-elicited defense in *Lotus* relies on a receptor pair consisting of the high-affinity, size-selective pseudokinase receptor CHIP13/14 and the low-affinity, active kinase co-receptor CERK6 and we have identified their functionally relevant chitin binding sites. To further define how these receptors interact during signaling, we conducted a Co-immunoprecipitation (Co-IP) assay with *Nicotiana benthamiana* leaves transiently expressing full-length CHIP13 and a kinase-inactive CERK6 in presence and absence of chitin (Fig. 4A). Notably, our Co-IP assay shows that the association of CERK6 with CHIP13 is greatly promoted in the presence of chitin, indicating a ligand-induced heterocomplex assembly (Fig. 4A, Exp. 1). This is confirmed by CHIP13 Y232A F233A, which does not recruit CERK6 upon chitin treatment (Fig 4A, Exp. 4). The Co-IP does not indicate that CERK6 forms a homocomplex *in planta*, neither in presence nor absence of chitin (Fig 4A, Exp. 3). In contrast, both wild-type CHIP13 and CHIP13 Y232A F233A form ligand-independent CHIP13-CHIP13 interactions (Fig. 4A, Exp. 2, 5). We similarly observe that the CHIP13 ectodomain forms a dimeric population in an *in vitro* SEC assay (fig. S9A).

**Fig. 4.**
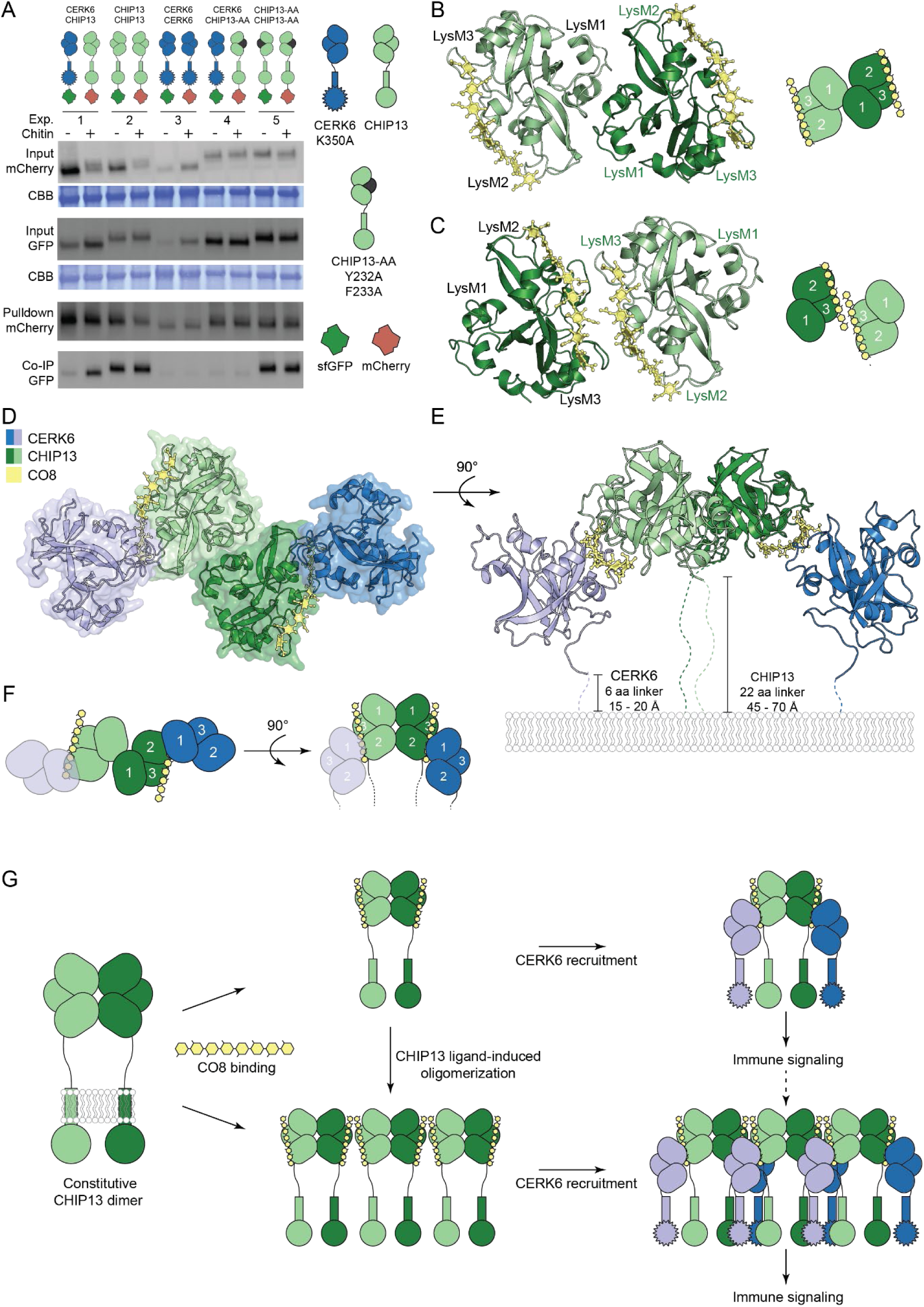
Model of the chitin perception complex in *Lotus japonicus*. (A) Co-immunoprecipitation experiments after transient expression of receptor-fluorophore fusion proteins in *N. benthamiana* leaves in the presence and absence of chitin. CBB indicates the Coomassie brilliant blue stained loading control. Exp. indicates different experimental receptor combinations assayed in presence/absence of chitin. (B) Ligand-independent CHIP13 dimer formation mediated by LysM1, bottom view. CHIP13 monomers are shown in light green and dark green cartoon representation, CO8 is drawn in yellow stick representation. Schematics indicate individual LysM domains and ligand placement. (C) Ligand-induced CHIP13 dimer, bottom view. Visualization and coloring scheme as in panel B. (D-F) Model of the tetrameric core chitin perception complex consisting of a LysM1-LysM1 CHIP13 dimer (light green and dark green) with chitin bound in LysM2 and LysM3. CERK6 (in light and dark blue) is recruited onto the reducing end of CO8 bound in CHIP13 LysM2. (D) Bottom view with surface and cartoon representations (E) 90° side view with schematic plasma membrane. Unmodelled linkers are indicated with dashed lines. (F) Schematic representation of panels D and E with numbering of individual LysM domains. (G) Model for perception of immunogenic long-chain chitin oligomers in *Lotus*.

We correlated the results of our Co-IP experiment with conserved interaction surfaces in our structural data. In all our five different crystal structures of CHIP13, LysM1 engages in the same ligand-independent contact with a symmetry-related LysM1 (Fig. 4B, fig. S9B-F, table S1). It is likely that the LysM1-LysM1 dimer reflects the constitutive CHIP13-CHIP13 interaction observed in Co-IP and SEC experiments. A second interface is found in all three CHIP13:CO8 crystal structures, where the reducing end of CO8 bound in LysM3 and the bridge domain form a ligand-induced dimer (Fig. 4C, fig. S9G-I). In crystal form 2 of CHIP13^33-274^:CO8 (PDB: 9Q83) the crystal packing almost exclusively consists of rows of alternating ligand-independent and ligand-induced interfaces (fig. S10A-B). This could indicate that CHIP13 not only forms ligand-independent complexes, but chitin potentially promotes CHIP13 oligomerization.

Using our ligand-bound CHIP13 and CERK6 structures we next modelled a chitin-induced heterocomplex. NAG1-3 and NAG6-8 of CHIP13-bound CO8 are accessible for a sandwich-like interaction as known from other LysM proteins (*20*, *21*, *25*, *26*). *In silico* docking CERK6 LysM1 onto the CHIP13 LysM3-bound CO8 reducing end (NAG1-3) causes non-resolvable sterical clashes with the N-glycan on CERK6 N46. This N-glycan, a conserved part of the LysM1 legume chitin signature motif, is predicted to be a high-confidence glycosylation site, and is present in all our crystal structures of CERK6 (table S1,(*5*). Additionally, CERK6 binding in CHIP13 LysM3 is incompatible with potential ligand-induced CHIP13 oligomerization, which is mediated by the reducing end NAGs. Contrary, similar binding onto the exposed non-reducing end in CHIP13 LysM2 (NAG 6-8) results in a sterically meaningful ligand-induced complex (fig. S11A). This arrangement is spatially sound due to the placement of CERK6 ‘below’ CHIP13 closer to the plasma membrane, fitting well with both length and alignment of the C-terminal linkers connecting the ectodomains to their transmembrane helices (Fig. 4D-F). Therefore, we propose that ligand-induced complex formation is mediated by CERK6 LysM1 associating onto the fixed, nonreducing end of CO8 in CHIP13 LysM2.

Our results provide evidence that CHIP13 forms a ligand-independent dimeric scaffold mediated by the LysM1-LysM1 interface and potentially further oligomerizes upon chitin binding with a ligand-induced mechanism. Chitin-bound CHIP13 homocomplexes recruit monomeric CERK6, triggering immune signaling. We propose a minimal tetrameric core receptor complex is sufficient to initiate signaling, while potential ligand-induced CHIP13 oligomers may further potentiate immunity through enhanced CERK6 recruitment (Fig. 4G, fig. S10C,D).

## Discussion

Our study provides the structural basis for the specific perception of long-chain chitin oligomers in plant immunity signaling. We show size-selectivity for long chitin oligomers is a hallmark of the CHIP-type LysM-RKs family and is facilitated through cooperative binding in the LysM2, LysM3, and the bridge domain. In contrast, previously characterized chitin receptors, e.g. *At*CERK1, *Os*CEBiP, *Lj*LYS11 or *Mp*LYR bind short chitins (CO4 or CO5) efficiently and affinity only increases slightly with increasing chitin lengths (*13*, *14*, *16*, *20*). Prediction models of CHIP-type LysM-RKs from different plant species reveal structural conservation to other established and candidate chitin receptors (fig. S3B(*9*, *11*, *27*). We propose long-chain chitin binding is similar throughout homologs to what we observe in our CHIP13:CO8 crystal structures. However, in some species CHIP-types may fulfill different functions, as e.g. rice chitin immunity appears to be mediated by CEBiP-type LysM-RPs (*24*)

Guided by our crystal structures, we identify a CH-π network as critical for the high chitin affinity in CHIP13 and required for LYR4 functionality in *Medicago*. Stacking and non-stacking CH-π interactions of aromatic side chains to pyranose and furanose rings are a well-known feature of other carbohydrate-binding proteins, such as carbohydrate binding modules, lectins and carbohydrate transporters, but have not been described for LysM-RK chitin binding before (*28*, *29*). Targeting the CH-π with Nb-aCHIP13 blocks signaling from the redundant receptors CHIP13/14 in wild-type *Lotus* seedlings and confirms their role in *Lotus* chitin immunity. Remarkably, our experiments show that targeted manipulation of plant pattern recognition receptor signaling through external application of specific nanobodies is possible in plant cells surrounded by a cell wall matrix.

Previously, we reported micromolar chitin affinities for CERK6 measured by microscale thermophoresis (*3*). We now conducted a more extensive, comparative binding analysis using ITC, STD-NMR, and Affinity PAGE that shows millimolar chitin affinity for both CERK6 and *Mt*CERK1. Despite its weak affinity, CERK6 binds CO5 in a sandwich-type mode using both LysM1 and LysM2, comparable to *Os*CERK1 (*20*). We have previously established LysM1, not LysM2, to be essential for legume CERK signaling, and validate our past prediction that the two conserved legume signature motifs in LysM1 indeed bind chitin (*5*). Their functional importance in signaling is further confirmed by specifically blocking LysM1 chitin binding using Nb-aCERK6 in wild-type *Lotus* seedlings. In contrast, the functionality of *Os*CERK1 and *At*CERK1 depends on LysM2 (*16*, *20*).

*At*CERK1 has high chitin affinity without size selectivity and was proposed to work either in a functional homocomplex or in heterocomplex with the CHIP-type receptors *At*LYK4 and *At*LYK5 (*9*, *16*, *25*, *30*, *31*). In many plant species engaging in AMS, CERK receptors have dual functions. In *Medicago*, *Mt*CERK1, together with *Mt*LYK8 (also a CERK-type receptor) are both required for establishment of AMS, while *lyr4* is unaffected in mycorrhizal colonization (*4*, *7*). Likewise, *Os*CERK1 acts in both AMS and immunity, where it is essential for symbiotic short-chain chitin oligomer signaling (*32*, *33*). The non-size selective high-affinity chitin binding of *At*CERK1 could reflect that *Arabidopsis* does not engage in AMS and has no strict requirement for stringent distinction between short and long chitin oligomers. In *Marchantia*, a single low-affinity/high-affinity receptor pair, *Mpa*CERK1/*Mpa*LYR, mediates both immunity and AMS using a non size-selective dosage-dependent mechanism, possibly indicating that vascular plants evolved more specialized receptors to fine-tune perception of symbiotic and pathogenic signals (*13*).

Together with previous studies, our findings support legumes and monocots employing low-affinity CERK-type coreceptors working in concert with high-affinity LysM-RKs or LysM-RPs. Similarly, leucine-rich repeat (LRR) peptide receptors in immunity or hormone signaling form high-affinity receptor/low-affinity co-receptor complexes, such as FLS2-BAK1 or HAESA-SERK1 (*34*, *35*). The LysM-RK pairs mediating root nodule symbiosis in *Lotus* and *Medicago*, NFR1/NFR5 and LYK3/NFP, however, all have high affinity for their lipochitooligosaccharide Nod factor ligand. This is possibly reflecting the inherent asymmetry and heterogeneity of Nod factors compared to chitin, a uniform molecule (*5*, *14*, *36*).

Our Co-IP and SEC experiments reveal that CHIP13 forms at least constitutive dimers. This finding is supported by our five CHIP13 crystal structures, each showing the LysM1-LysM1 ligand-independent interface (Fig. 4B, fig. S9). CHIP13 can therefore be placed among single-pass receptor kinases forming constitutive dimers/oligomers such as *Arabidopsis* LRR receptor kinases FLS2 and SUB and LysM-RKs *At*LYK5, AtLYK4 and *Lj*NFR5 (*11*, *37–39*). We do not observe a CERK6 homocomplex in our pull-down assay, suggesting the CERK6 ligand-induced LysM1-LysM2 dimer seen *in crystallo* is not physiologically relevant.

We find an additional, ligand-induced interface in all our CHIP13:CO8 structures, opening the possibility for further ligand-induced oligomerization of CHIP13 (Fig. 4C, fig. S10). The ligand-induced CHIP13 oligomer scaffold has the correct ectodomain alignment and is spatially compatible with our proposed position for CERK6 onto the nonreducing end of CO8, (fig. S10). Chitin treatment transcriptionally upregulates *Chip13/14* in *Lotus* roots (*22*), and we speculate that the continuing presence of chitin could promote the formation of larger receptor scaffolds facilitating amplified chitin immunity signaling.

Previous studies suggested differing models on how LysM-RK and LysM-RP complexes are formed to initiate signaling (*11*, *16*, *25*, *30*, *40*). Based on our structural, biochemical and functional data, we provide a mechanistic model, where the size-selective, high-affinity CHIP13 homodimer acts as a constitutive scaffold, recruiting two monomeric, low-affinity CERK6 receptors into a ligand-induced, signaling-competent tetrameric core complex.

## Supporting information

Table S1. Data collection and refinement statistics.

## Acknowledgements

The authors thank Thi Bich Luu for providing *Lotus japonicus cerk6* seeds, Bine Wissendorf Simonsen for support with *Medicago* plant experiments, Majken Kiel Sørensen for greenhouse work and Simona Radutoiu for helpful discussions. The synchrotron data for 9GXF, 9H3A, 9Q84, 9Q83 and 9QRS was collected at beamlines P13/P14 operated by EMBL Hamburg at the PETRA III storage ring (DESY, Hamburg, Germany) under proposals MX-377 and MX-846. We thank Johanna Hakanpää and Isabel Bento for assistance in using the beamline. Synchrotron data for 9GXZ, 9H39, 9H3B, 9H24 and 96HV was collected at MAX IV. We acknowledge MAX IV Laboratory for time on Beamline BioMAX under proposal 20220062. Research conducted at MAX IV, a Swedish national user facility, is supported by the Swedish Research council under contract 2018-07152, the Swedish Governmental Agency for Innovation Systems under contract 2018-04969, and Formas under contract 2019-02496. We thank Ana Gonzalez and Mirko Milas for assistance in using the beamline. This work was supported by an instrument grant for a PEAQ-ITC from the Carlsberg Foundation (CF22-0971) to Esben Lorentzen. We thank the Nanobio ICMG (UAR 2607) mass spectrometry platform for technical support and acknowledge funding by Agence Nationale de la Recherche through Labex ARCANE and CBH-EUR-GS (ANR-17-EURE-0003), Glyco@Alps (ANR-15-IDEX-02). This work was funded by the Novo Nordisk Foundation (NNF18OC0052855), the Danish Council for Independent Research (3103-00137B), the Carlsberg Foundation (CF21-0139) and the Danish National Research Foundation (DNRF79).

## Author contributions

KG: Conceptualization, study design, project supervision, all crystal structures, structural analysis and visualization, molecular modeling and docking, protein production, biochemistry, *in planta* nanobody assays, nanobody work. SBH: crystal structures of *Mt*CERK1, CERK6:CO5, and CERK6:Nb, biochemistry, protein production, nanobody work. HR: crystal structure of LYR4, biochemistry, *Medicago* complementation, Co-IP. HA: nanobody work, protein production, biochemistry, and crystal structure of CERK6:Nb. EM: NMR, nanobody work. JC: Crystal structure of CERK6. CB: biochemistry. ECP: biochemistry. MV: protein production. SF: ligand production. JS: project supervision. KRA: project supervision, study design, nanobody work. KG, SBH, HR and KRA prepared the manuscript with input from all authors.

## Data availability

All data are available in the main text or the supplementary materials. Coordinates and structure factors for all crystal structures are deposited in the Protein Data Bank CHIP13: 9GXF; CHIP13:CO8: 9H3A; CHIP13:Nb-aCHIP13: 9H39; CHIP13^33-274^:CO8 crystal form 1: 9Q84; CHIP13 (33–274):CO8 crystal form 2: 9Q83; *Mt*LYR4: 9GXZ, CERK6:CO5: 9H3B, CERK6:Nb-aCERK6-1:Nb-aCERK6-2: 96HV, CERK6: 9QRS, *Mt*CERK1: 9H24

## Supplementary Materials

Materials and Methods

Figs. S1 to S11

Tables S1 to S4

