## Supplementary Materials for "Structural basis for size-selective perception of chitin in plants"

### Materials and Methods

LysM receptor ectodomain expression and purification

*Lotus japonicus* CHIP13$^{33\text{-}253}$, CHIP13$^{33\text{-}274}$, LYR4$^{24\text{-}242}$ and *Mt*CERK1$^{30\text{-}232}$ were codon-optimized for insect cell expression (Genscript, Piscataway, United States of America) and cloned into the pOET4 baculovirus transfer plasmid (Oxford Expression Technologies) using *XhoI* and *HindIII* restriction sites. Proteins were targeted for secretion using the gp64 signal peptide from the *Autographa californica* nucleopolyhedrovirus (AcMNPV). CHIP13$^{33\text{-}253}$ Y232A F233A was prepared by site-directed mutagenesis from the wild-type construct using the Q5® Site-Directed Mutagenesis Kit (New England Biolabs).

Recombinant baculoviruses for protein expression were produced using the FlashBac Gold kit (Oxford expression technologies) according to the manufacturer's instructions in *Spodoptera frugiperda* Sf9 cells with lipofectin (Thermo Fisher Scientific) or BaculoFECTIN (Oxford Expression Technologies) as transfection reagent. Sf9 cells were cultured in suspension in serum-free HyClone SFX medium (Cytiva) supplemented with 1% antibiotic-antimycotic (Gibco) and 1% CD lipid concentrate (Gibco). Expression was induced by addition of the respective passage 3 virus at a cell density of $10^6$ cells/ml. After 5 days, the medium was cleared by centrifugation and dialyzed overnight against 50 mM Tris-HCl pH 8, 200 mM NaCl at 4°C. Recombinant ectodomains were captured by two rounds of Ni-IMAC purification, first with a HisTrap excel column, then with a HisTrap HP column (both Cytiva). As a final purification and quality control step, ectodomains were purified by size exclusion chromatography on Superdex 75 Increase 10/300 and Superdex 200 Increase 10/300 columns (Cytiva) in either phosphate buffered saline, pH 7.2 (for binding assays) or 20 mM HEPES pH 7.5, 150 mM NaCl (crystallization buffer).

*At*CERK1 and CERK6 ectodomains were produced as previously described (*3*, *14*, *41*).

Nanobody selection

The procedure was conducted similarly as previously described (*38*, *42*). A llama (*Lama glama*) was immunized and boosted four times with 100 µg of each purified CHIP13 and CERK6 LysM ectodomains. Peripheral blood lymphocytes were isolated from a blood sample, and RNA was extracted using the RNase Plus Mini Kit (Qiagen). cDNA was synthesized using the Superscript III First Strand Kit (Invitrogen) with random hexamer and dTTTT primers. The nanobody (Nb) coding regions were amplified via several tandem PCRs and inserted into a phagemid vector, fusing the Nbs C-terminal to an E-tag and the phage pIII coat protein. The M13 phage display library was generated using VCSM13 helper phage. LysM ectodomains were biotinylated for selection using the Chromalink NHS labeling system (Solulink). Biotinylated antigen (20 µg) was added to 100 µl of MyOne Streptavidin T1 Dynabeads (Thermo Fisher Scientific) in PBS containing 1x fish gelatin (Biotium). M13 phage particles ($2.5 \times 10^{13}$) were incubated with antigen-coated Dynabeads for 1 hour, followed by 15 washes with 1 ml of PBS containing 0.1% Tween-20. Phages were eluted by incubating the beads in 0.2 M glycine (pH 2.2) for 15 minutes, and the elution was immediately neutralized by adding 1 M Tris buffer (pH 9.1) to restore a neutral pH. The eluted phages were amplified and subjected to a second round of phage display with reduced amounts of biotinylated antigens (2 µg) and M13 phage particles ($2.5 \times 10^{11}$). After two selection rounds, single colonies were grown in LB medium in 96-well plates for 6 hours. Nb expression was induced with 1 mM IPTG and incubated overnight at 30°C and 150 rpm. The plates were centrifuged, and 50 µl of supernatants were transferred to ELISA plates coated with 0.1 µg of LysM ectodomain antigens and blocked with PBS containing 0.1% Tween-20 (PBS-Tween) and 1x fish gelatin. Supernatants were incubated for 1 hour, then washed six times with PBS-Tween and probed with anti-VHH-HRP antibody (Bethyl) diluted 1:10,000 in PBS-fish gelatin buffer, followed by three rounds of wash. The plates were developed with TMB substrate, quenched with 1 M HCl, and absorbance was measured at 450 nm. In parallel, mock selections and ELISA validations were performed to identify and exclude false positives. Phagemids from positive clones were sequenced, and Nb-encoding DNAs were cloned into pET22b(+) (Novagen) for bacterial expression.

Nb-aCERK6-1 was generated by grafting the complementary-determining regions of the selected nanobody into the scaffold of the anti-transcobalamin II nanobody 4 (TC-Nb4) (*43*).

<u>Expression and purification of nanobodies</u>

Specific nanobodies against CERK6 and CHIP13 ectodomains were recombinantly expressed in *Escherichia coli* LOBSTR or BL21 Rosetta2 strains. Transformed cells were induced with 0.2 mM Isopropyl β-D-1-thiogalactopyranoside (Sigma-Aldrich) after reaching an OD600 = 0.5 and incubated shaking at 18°C for 16 hours. Cells were harvested by centrifugation, resuspended in Lysis buffer (50 mM Tris-HCl pH 8, 500 mM NaCl) and subsequently lysed by sonication. Nanobodies were captured by affinity purification using a HisTrap HP affinity column (Cytiva) and eluted with a step gradient using elution buffer (50 mM Tris-HCl pH 8, 200 mM HCl, 250 mM Imidazole). As a final purification step, nanobodies were gel filtrated on a Superdex 75 16/600 prep grade column on an ÄKTA pure system (Cytiva). For crystallography, nanobodies were gel filtrated into crystallization buffer (20 mM HEPES, pH 7, 150 mM NaCl). For nanobody assays on *Lotus* seedlings, plant assay buffer was used during SEC (10 mM MES, pH 6.0, 50 mM KCl).

<u>Crystallization and Structure Determination</u>

All crystallographic data were collected at either EMBL beamlines P13/P14 (DESY, Hamburg, Germany) or BioMax (MaxIV, Lund, Sweden) as indicated in the specific paragraphs and Table S1 (*44*, *45*). Data processing and reduction for all datasets was done using the XDS package unless stated otherwise (*46*). Molecular replacement was conducted in phenix.phaser using the specified search models below (*47*). All crystal structures were built in COOT (*48*) and refined using phenix.refine from the phenix suite (*49*). Geometry was assessed in Molprobity (*50*).

Crystals for **CHIP13** without ligand (PDB 9GXF) were obtained in a sitting drop vapor diffusion setup in 0.1 M sodium acetate trihydrate pH 4.5, 23% PEG-3350 at 19°C and a protein concentration of 10 mg/ml. Crystals were cryoprotected in sodium acetate trihydrate pH 4.5, 23% PEG-3350, 10 % (v/v) glycerol before snap-cooling in $LN_2$. Diffraction data were collected at the DESY P14 beamline to d = 1.64 Å. Phases were obtained by molecular replacement (MR) with a predicted model of CHIP13 ectodomain without its N-terminal bridge domain (aa 56-242) generated in Colabfold as search model (*51*, *52*). The trimming of low confidence regions (LDDT < 0.7) and conversion of LDDT values was done with phenix.process_predicted_model (*49*, *53*). This structure served as MR search model for all other structures of CHIP13 and LYR4.
Crystals of the **CHIP13:CO8** complex **(**PDB: 9H3A**)** were grown in a sitting drop vapor diffusion setup in 0.2 M ammonium chloride, 20 % (w/v) PEG-3350. Cryoprotection was done in 0.2 M ammonium chloride, 20 % w/v PEG-3350, 10% (v/v) ethylene glycol before snap-cooling in $LN_2$. Diffraction data to d = 1.35 Å were collected at the DESY P14 beamline. Non-water hydrogens were modeled explicitly due to high resolution.

Crystals of the **CHIP13 (33-274):CO8** complex were obtained in two different crystal forms (PDB: 9Q83 and 9Q84). The first (PDB: 9Q84) was produced in sitting drop vapor diffusion in 0.2 M NaCl, 0.1 M Bis-Tris pH 5.5, 25% (w/v) PEG-3350 and cryoprotected in 0.2 M NaCl, 0.1 M Bis-Tris pH 5.5, 25% (w/v) PEG-3350, 10% (v/v) glycerol. Diffraction data to d = 1.4 Å were obtained at the DESY P14 beamline. In this high-resolution structure, non-water hydrogens were explicitly modeled. The second crystal form (PDB: 9Q83) was obtained in 0.15 M LiSO4, 0.075 M sodium acetate pH 4.5 22.5% PEG-8000 and cryoprotected in  0.15 M LiSO4, 0.075 M sodium acetate pH 4.5 22.5% PEG-8000, 10% (v/v) glycerol. Diffraction data were obtained at the DESY P13 beam-line. Due to anisotropy, diffraction data of crystal form 2 were further processed by anisotropic scaling using the STARANISO server (*54*) along an ellipsoidal diffraction limit surface (1.99 Å, 2.90 Å, 2.57 Å) defined by the following criteria: $R_{pim}$ <= 0.6, I/sig(I) >= 2.0, CC(1/2) >= 0.3.

Crystals of CHIP13 in complex with CO8 were obtained by co-crystallization. Powdered ligand (*55*) was directly added to 10 mg/ml of the respective CHIP13 protein to ligand saturation (esti-mated 1-2 mM). The complex was incubated for two hours at room temperature before clearing by centrifugation and setting up crystallization experiments.

Crystallization of **CHIP13:Nb-aCHIP13** (PDB: 9H39) complex was achieved by mixing purified CHIP13 and Nb-aCHIP13 in a 1:1.1 molar ratio and purifying the complex on gel filtration in crys-tallization buffer on a Superdex 75 increase 10/300 column (Cytiva). The purified complex was crystallized in a sitting drop vapor diffusion setup in 0.2 M sodium acetate 0.1 M Bis-Tris propane, pH 7.5 20 % (w/v) PEG-3350 at 5.5 mg/ml and cryoprotected in in 0.2 M sodium acetate 0.1 M Bis-Tris propane, pH 7.5 20 % (w/v) PEG-3350, 10% (v/v) ethylene glycol. Diffraction data to d=1.66 Å were collected at BioMax. The phase problem was solved by MR with the structure of CHIP13 (PDB: 9GXF) and the nanobody scaffold from PDB: 6QUP, chain B, as search models (*42*).

Crystallization of **LYR4** (24-242**,** PDB 9GXZ**)** was facilitated by deglycosylation of the protein with a mix of the endoglycosidases Endo F1 and Endo F3 overnight at room temperature. The gly-can-trimmed protein was then SEC purified in crystallization buffer. Crystals were grown in a sit-ting drop vapor diffusion setup in 0.1 M Bis-Tris pH 5.5, 2.0 M ammonium sulphate at 9.1 mg/ml and cryoprotected in 0.1 M Bis-Tris pH 5.5, 2.0 M ammonium sulphate, 25% (v/v) ethylene glycol before snap-cooling in $LN_2$. Diffraction data to d = 1.50 Å were collected at BioMax.

The crystal structure of **CERK6:CO5** (PDB: 9H3B) was obtained by direct addition of powdered chitopentaose (Megazymes) to 6.5 mg/ml purified protein to a final concentration of 5 mM carbo-hydrate. Crystals were obtained in a sitting drop vapor diffusion setup in 0.2M Potassium sodium tartrate tetrahydrate, 0.1M Bis-Tris propane pH 6.5, 18% (w/v) PEG-3350. Crystals were cryopro-tected in 0.1M Bis-Tris propane pH 6.5, 18% (w/v) PEG-3350, 10 % (v/v) PEG-200 before snap-cooling in $LN_2$. Diffraction data were collected at DESY P14 and were further processed by aniso-tropic scaling using the STARANISO (*54*) server  along an ellipsoidal diffraction limit surface (4.09 Å, 4.09 Å, 2.43 Å) defined by the following criteria: $R_{pim}$ <=  0.6, I/sig(I) >= 2.0, CC(1/2) >= 0.3. The phase problem was solved by MR using CERK6 (PDB: 5LS2) as search model (*3*).

Crystallization of the **CERK6:aCERK6-Nb:aCERK6-Nb2** (PDB: 96HV) complex was achieved by purification of the complex by SEC in crystallization buffer, similar to the CHIP13:Nb-aCHIP13 complex. Crystals were obtained in sitting drop vapor diffusion setup at 19° C in a condition containing 0.2 M sodium bromide, 20 % (w/v) PEG 3350 and cryoprotected in 10 % (v/v) ethylene glycol. Diffraction data were collected at the BioMax beamline. Due to the low-symmetry triclinic space group, two individual datasets were collected from one single crystal at an offset of κ = 30°. After standard data processing in XDS, datasets were scaled relative to one another in XSCALE (*46*), resulting in a single dataset with increased completeness. Diffraction data were further processed by anisotropic scaling using the STARANISO server (*54*) along an ellipsoidal diffraction limit surface (2.19 Å, 3.34 Å, 2.43 Å) defined by the following criteria: $R_{pim}$ <= 0.6, I/sig(I) >= 2.0, CC(1/2) >= 0.3. The resulting dataset has a resolution of d = 3.05 Å anisotropically extended to 2.01 Å. The phase problem was solved using CERK6 (PDB:5LS2) and the nanobody scaffold from PDB:6QUP, chain B as search models (*3*, *42*).

Improved crystals of **CERK6** (PDB 9QRS) were obtained in a sitting drop vapor diffusion setup at 4°C at 5 mg/ml in a condition containing 0.2 M ammonium sulphate, 0.1 M sodium acetate pH 4.6 and 30% (w/v) PEG-2000 MME. Before snap-cooling in $LN_2$, crystals were cryoprotected in 0.1 M sodium acetate pH 4.6, 30% (w/v) PEG-2000 MME and 10% (v/v) ethylene glycol. Diffraction data to d = 1.87 Å were collected at the DESY P13 beamline.
***Mt*CERK1** (PDB: 9H24) was crystallized at 19° C in sitting drop vapor diffusion in a condition containing 0.2 M Lithium sulfate, 0.1 M Tris 8.5, 40 % v/v PEG 400. Diffraction data to d = 2.98 Å were collected at BioMax and processed using the EDNA pipeline on the beamline to d = 3.0 Å (*56*). CERK6 (PDB: 5LS2) was used as MR search model (*3*).

All visualizations of structural models were prepared using PyMol 3.1.1.

Structural Modeling of Receptor-ligand complexes
Homology modeling was performed with Alphafold 2 provided by Colabfold v. 1.3.0. using the MMseqs2 database in inpaired+paired mode without PDB templates (*51*, *52*). Protein structures were predicted using the ptm model type with 3 recycling per model. The model with the highest overall pLDDT (out of 5) was chosen for further analysis.

Isothermal titration calorimetry (ITC)

*In vitro* binding of ectodomains to chitin oligomers was measured on an PEAQ-ITC isothermal titration calorimetry system (Malvern) at 25°C in a PBS buffer (pH 7.4, Sigma) with receptor ectodomain in the microcell and chitin ligand in the syringe, respectively (CO5: Megazymes, CO8 (*55*). Concentrations are shown in the table below. An experiment consisted of 20 injections in total, the first with 0.4 µl and 19 2 µl injections with 150 sec spacing between injections. Corresponding control experiments with buffer instead of receptor were conducted. ITC data were analyzed in MicroCal PEAQ-ITC Analysis software and plotted in Graphpad Prism.

| Experiment | Microcell concentration [mM] | Syringe concentration [mM] |
|---|---|---|
| CHIP13 – CO8 | 0.025 | 0.5 |
| CHIP13 – CO5 | 0.1 | 10 |
| CHIP13:Nb-aCHIP13 – CO8 | 0.025 | 1 |
| CHIP13 Y232A F233A – CO8 | 0.05 | 1 |
| CHIP13 Y232A F233A – CO5 | 0.08 | 10 |

| | | |
|---|---|---|
| CHIP13 $^{33\text{-}274}$ – CO8 | 0.03 | 0.3 |
| CHIP13 $^{33\text{-}274}$ – CO5 | 0.05 | 10 |
| MtLYR4 – CO8 | 0.05 | 1 |
| MtLYR4 – CO5 | 0.1 | 5 |
| CERK6 – CO8 | 0.25 | 1 |
| CERK6 – CO5 | 0.25 | 15 |
| MtCERK1 – CO8 | 0.1 | 1 |
| MtCERK1 – CO5 | 0.25 | 10 |

Biolayer interferometry (BLI)

Direct binding of receptor ectodomains to nanobodies was measured on an OctetRED 96 biolayer interferometry system (Sartorius). Purified CHIP13 and CERK6 were biotinylated using the Chromalink Biotin Labeling System (Solulink) by NHS-biotin chemical linking of primary amines on the protein surface. First, ectodomains were buffer exchanged on SEC to phosphate-buffered saline, pH 7.4 (Sigma). 330 µg protein was incubated for 90 minutes at room temperature with NHS-biotin dissolved in dimethylformamide at 10 mg/mL. The required amounts of NHS-biotin were determined according to the manufacturer's instructions. After incubation, a second round of SEC was performed to remove excess biotin.
Biotinylation levels of the final protein solutions was estimated by measuring absorbance at λ = 280 nm and λ = 354 nm on a UV–Vis NanoDrop ND-1000 spectrophotometer (Thermo Scientific) and using the manufacturer's Chromalink label calculator. Each protein was estimated to contain approximately 3–4 biotin molecules per protein molecule. Binding experiments were performed in phosphate buffered saline pH 7.4, 0.01% Tween-20 at 25°C under 1000 rpm agitation using black polystyrene 96-well plates (Sarstedt). Biotinylated ectodomains were immobilized on streptavidin biosensors (Kinetic Quality, Sartorius ForteBio) to an immobilization level of 0.5 nm. After a 60 second wash step, association was measured for 600 seconds and dissociation for 300 seconds. Nanobodies were titrated in serial 1:2 dilutions in the indicated concentrations.
Binding data were analyzed using the ForteBio Data analysis software and GraphPad Prism (Version 10.4.1, Graphpad Software LLC). Fits were performed by non-linear regression (Global fit to association then dissociation model with shared $k_{on}$, $k_{off}$ and $B_{max}$) values.

Affinity PAGE (APAGE)

Shrimp-shell chitin (Sigma-Aldrich) was homogenized in Tris-HCl pH 8.8 using a pestle head mounted on an electrical drill for 5 min before being mixed into the separation gel mixture at 0.25 % (w/v). Affinity gels were cast with a 5% acrylamide stacking gel, pH 6.8 without chitin and 5 µg protein sample was loaded in each lane. A control native PAGE was performed in parallel without addition of shrimp-shell chitin. Gels were run in a 25 mM Tris/glycine pH 8.3 running buffer system in parallel at 4 °C for 5 h with a constant voltage of 150 V.

STD-NMR

All samples were prepared in phosphate buffered saline in $D_2O$, pH 7.4 by buffer exchange of purified protein in a spin column. The following protein:ligand concentrations were tested: 20 µM:1000 µM (1:50), 40 µM:1000 µM (1:25) and 60 µM: 300 µM (1:5). All spectra were acquired at 25 °C on a Bruker spectrometer (500 MHz) equipped with a 1H-13C-15N 5 mm TXI liquid-state probe. Saturation transfer difference (STD) experiments were set up following (*57*) using the STD pulse program (*58*, *59*).The shaped pulse was calibrated with a selzg experiment optimizing for p11, and SPNAM1 was defined as gauss1.1000 for the protein saturation. The on-resonance irradiation of the protein was set in the aliphatic region (0.50, 0.57, 0.60, 0.80 ppm), and the off-resonance frequency at -30 ppm. The saturation time (d20) was 3s.

Generation of plant expression vectors

For the generation of *Medicago truncatula* expression vectors, *Lyr4* constructs were assembled with the *LYR4* promoter, *35S* terminator and a 6xHistidine tag in the pIV10_tYFP-NLS expression vector by Golden Gate cloning (*60*, *61*). YFP fused to a nuclear localization sequence (NLS) served as a transformation control. For immunoprecipitation experiments in *Nicotiana benthamiana*, *Chip13/Lys13* and *Cerk6* constructs were assembled with a *Ubiquitin* or *35S* promoter, a *35S* or *NOS* terminator and a GFP- or mCherry-tag in the pICH binary vector backbone (*62*).

Hairy root transformation

*Medicago* R108 and *lyr4* seeds were scarified in sulphuric acid for 3 min and surface-sterilized with diluted bleach solution (3%, 4 minutes). Seedlings were germinated on wet filter paper (AGF 651; Frisenette ApS) in sterile Petri dishes at 21°C for two days and then transferred to agar plates (0.8% Gelrite, Duchefa Biochemie) supplemented with ½ Gamborg's B5 nutrient solution (Sigma-Aldrich). Transconjugant *Agrobacterium rhizogenes* AR1193 strains carrying the construct of interest were grown for three days on LB Agar containing 100 µg/mL Ampicillin, Rifampicin and Spectinomycin and cells were resuspended in YMB (5 g/L mannitol, 0.5 g/L yeast extract, 0.5 g/L $K_2HPO_4$, 0.2 g/L $MgSO_4$ x 7 $H_2O$, 0.1 g NaCl, pH = 6.8). 5-day-old seedlings were transformed with the bacterial suspension using a 1 mL syringe with a needle (Sterican Ø 0.40x20 mm), punching the hypocotyl and placing a droplet on the wound. The transformed seedlings were left in the dark for one day and then moved to 21°C under 16/8-h light/dark conditions. After three weeks, primary roots were cut off and seedlings with transformed roots were moved to Magenta boxes (Sigma-Aldrich) filled with clay aggregate (LECA, 2-4 mm; Saint-Gobain Weber A/S) and 80 mL ¼ B&D nutrient solution supplemented with 3 mM $KNO_3$.

*Medicago* oxidative burst experiments

*Lyr4* constructs were tested for their ability to complement the *Medicago lyr4* mutant in reactive-oxygen species production in response to chitin. Three-week-old hairy roots from individual plants producing the tYFP-NLS transformation marker were cut into 1 mm pieces and similar amounts of root material were transferred into 96-well flat-bottomed polystyrene plates (Greiner Bio-One) with 180 µL of water in each well. After overnight incubation at room temperature, the water was replaced by a reaction mixture consisting of 0.5 mM L-012 (Wako Chemicals), 5 µg/mL horseradish peroxidase (Sigma-Aldrich), and 1 µM octa-N-acetyl-chitooctaose (CO8). Luminescence was recorded with a Varioskan LUX multimode microplate reader (Thermo Fisher Scientific) for 30 minutes.

Nanobody treatment of *Lotus* seedlings and *Lotus* oxidative burst experiments

*Lotus japonicus* (ecotype Gifu) and *cerk6* seeds were scarified with sandpaper and surface-sterilized with diluted bleach solution (1%, 15 minutes). Seedlings were germinated on wet filterpaper (AGF 651; Frisenette ApS) in sterile Petri dishes at 21°C for 60 hours. Nanobodies were concentrated in centrifugal columns (Sartorius Stedim, 5 kDa MWCO) to 250 µM in plant assay buffer (10 mM MES pH 6.0, 50 mM KCl) before dilution in sterile MilliQ water to the final assay concentration of 5 µM nanobody. Single seedlings were transferred to 96-well flat-bottomed polystyrene plates (Greiner Bio-One) and treated with either 180 µl purified nanobody solution or water. After 4-6 hours of incubation at room temperature in the dark, the solution was replaced by a reaction mixture consisting of 0.5 mM L-012 (Wako Chemicals), 5 µg/mL horseradish peroxidase (Sigma-Aldrich), and 1 µM octa-N-acetyl-chitooctaose (*55*). Luminescence was recorded with a Varioskan LUX multimode microplate reader (ThermoFisherScientific) for 30 minutes.

*Agrobacterium* mediated transient transformation of *Nicotiana benthamiana* plants

Cultures of *Agrobacterium tumefaciens* AGL1 carrying the construct of interest were resuspended in infiltration solution (10 mM $MgCl_2$, 10 mM MES, 150 µM acetosyringone, pH 5.6) to an $OD_{600}$ = 0.25. An *A. tumefaciens* strain transformed with the RNA-silencing suppressor p19 was added to each solution to a final $OD_{600}$ = 0.025. The solutions were incubated at room temperature for two hours in the dark and infiltrated with a blunt end syringe into the abaxial side of *N. benthamiana* leaves from four- to five-week-old plants. Two days after infiltration, leaves were harvested.

Co-immunoprecipitation assay

*N. benthamiana* leaves were ground to powder using porcelain beads. Extraction buffer containing 50 mM Tris-HCl, pH 7.5, 150 mM NaCl, 10% (v/v) glycerol, 2 mM EDTA, 1 mM phenylmethyl-sulfonyl fluoride, Protease Inhibitor Cocktail (Sigma-Aldrich), and 1% (v/v) IGEPAL CA-630 (Alfa Aesar) was added to the powder. Samples were divided and ground shrimp-shell chitin (Sigma-Aldrich) was added to half of the samples at a final concentration of 0.5% w/v. After incubation for 2 hours at 4°C on a rotor, samples were centrifuged at 4,700 rpm, 30 min, 4°C and the supernatants were transferred to new tubes. Magnetic RFP beads (Chromotek) were added to the supernatant and the samples were incubated at 4°C overnight. Beads were collected using a magnetic rack and washed four times with extraction buffer. To release bound protein, 100 µL SDS sample buffer (25 mM Tris-HCl, pH 6.8, 10% (v/v) glycerol, 2% SDS, 0.2% 2-mercaptoethanol, 0.6 mg/mL Bromophenol blue) was added and samples were heated at 95°C for 5 min.

Western Blot

Proteins were separated on 12% SDS-PAGE gels at 200 V for 45 min and blotted to PVDF membranes (Merck) by wet transfer for 2 hours at 90 V. Membranes were blocked with 5% milk in PBS with 0.1% Tween-20 for 2 hours at room temperature. For detection of GFP, membranes were incubated overnight at 4°C with mouse anti-GFP (632381, TaKaRa, 1:5000) in 2.5% milk in PBS with 0.1% Tween-20 followed by anti-mouse HRP (A4416; Sigma-Aldrich, 1:10,000) for 2 h in 2.5% milk in PBS with 0.1% Tween-20. For detection of mCherry, membranes were incubated overnight at 4°C with rabbit anti-mCherry (632496, TaKaRa, 1:5000) in 2.5% milk in PBS with 0.1% Tween-20 followed by anti-rabbit HRP (A6154; Sigma-Aldrich, 1:10,000) for 2 h in 2.5% milk in PBS with 0.1% Tween-20. Chemiluminescence was detected on a G-BOX (Syngene) using ECL Prime Western Blotting Detection Reagent (Cytiva). Membranes with input samples were subsequently stained with Coomassie for the detection of total protein.

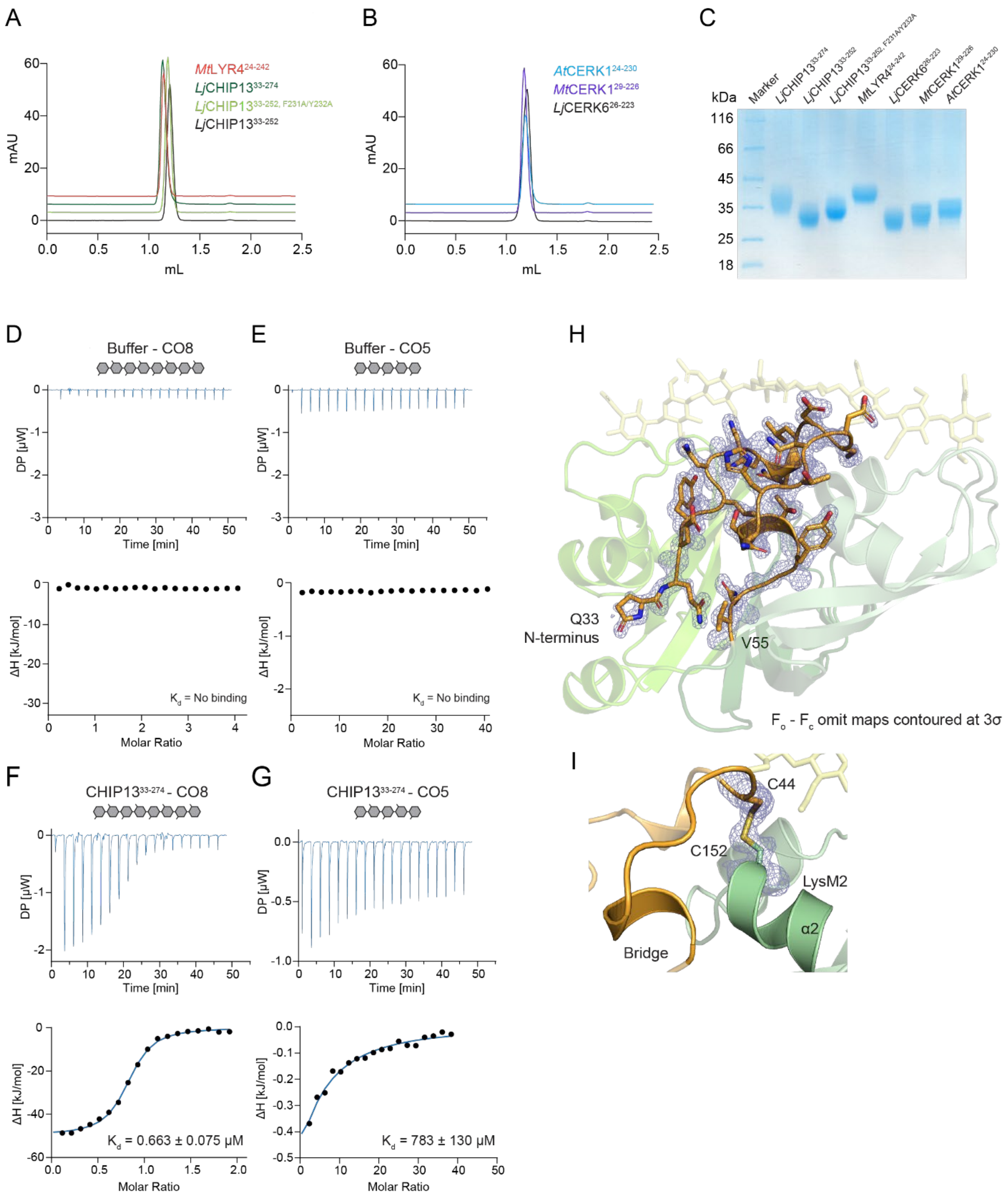

A
B
C
MtLYR4 24-242
LjCHIP13 33-274
LjCHIP13 33-252, F231A/Y232A
LjCHIP13 33-252
AtCERK1 24-230
MtCERK1 29-226
LjCERK6 26-223
mAU
mL
kDa
Marker
116
66
45
35
25
18
D
E
H
Buffer - CO8
Buffer - CO5
DP [μW]
Time [min]
ΔH [kJ/mol]
Molar Ratio
K<sub>d</sub> = No binding
Q33
N-terminus
V55
F<sub>o</sub> - F<sub>c</sub> omit maps contoured at 3σ
I
C44
C152
LysM2
α2
Bridge
F
G
CHIP13 33-274 - CO8
CHIP13 33-274 - CO5
K<sub>d</sub> = 0.663 ± 0.075 μM
K<sub>d</sub> = 783 ± 130 μM

**Fig. S1. Receptor ectodomains expressed and purified from insect cells and ITC control experiments.**

(A) $A_{280}$ chromatogram of SEC analysis of *Mt*LYR4$^{24\text{-}242}$, CHIP13$^{33\text{-}274}$, CHIP13$^{33\text{-}253}$, CHIP13$^{33\text{-}253}$ Y232A F233A on an analytical Superdex 75 increase, 3.2/300 column. (B) $A_{280}$ profiles of SEC runs of *At*CERK1$^{24\text{-}230}$, *Mt*CERK1$^{29\text{-}226}$, *Lj*CERK6$^{26\text{-}223}$ on an analytical Superdex 75 increase, 3.2/300 column. (C) SDS-PAGE analysis of ectodomains analyzed in (A) and (B). All proteins are glycosylated and migrate larger than their theoretical molecular weight. (D, E) Control ITC isotherms for titration of CO8 (D) and CO5 (E) into buffer. (F+G) ITC isotherms for titration of CO8 (F) and CO5 (G) into purified complete ectodomain of CHIP13 (CHIP13$^{33\text{-}274}$). (H) $F_o - F_c$ omit map of bridge domain (amino acids Q33-V55) electron density in the CHIP13:CO8 crystal structure (PDB: 9H3A) contoured at 3σ. Resolution is d = 1.35 Å. N-terminal pyroglutamate formation of Q33 is apparent. (I) $F_o - F_c$ omit map of additional disulphide C44:C152 contoured at 3σ. Resolution is d = 1.35 Å.

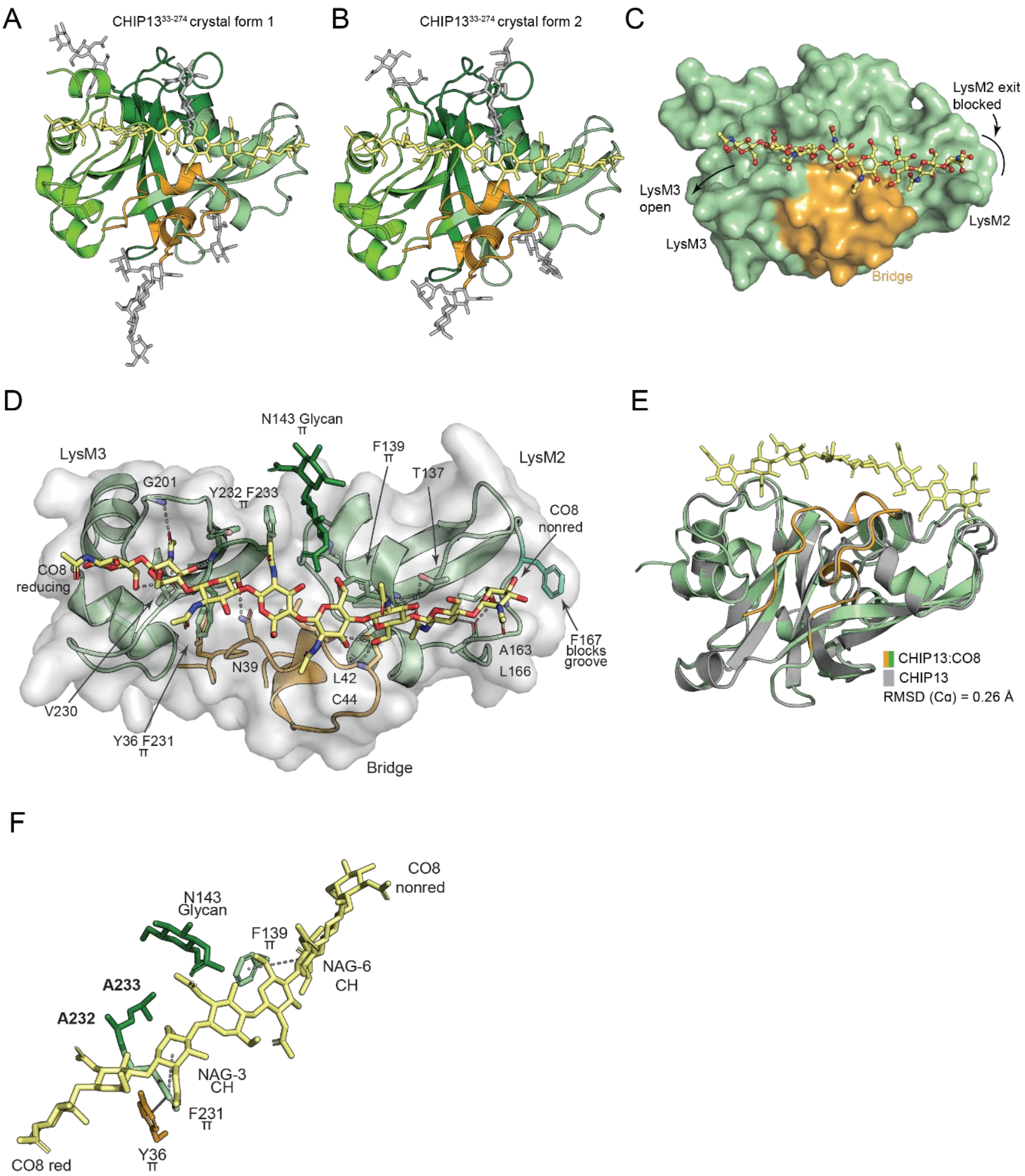

A
CHIP13^33-274 crystal form 1
B
CHIP13^33-274 crystal form 2
C
LysM2 exit
blocked
LysM3
open
LysM3
LysM2
Bridge
D
N143 Glycan
π
LysM3
F139
π
LysM2
T137
G201
Y232 F233
π
CO8
nonred
CO8
reducing
N39
L42
C44
A163
L166
F167
blocks
groove
V230
Y36 F231
π
Bridge
E
CHIP13:CO8
CHIP13
RMSD (Cα) = 0.26 Å
F
CO8
nonred
N143
Glycan
F139
π
NAG-6
CH
A233
A232
NAG-3
CH
F231
π
Y36
π
CO8 red

**Fig. S2. CHIP13 is a specific receptor for long chitins and uses CH- π interactions for high-affinity ligand binding.**
(A+B) Crystal structures of the CHIP13$^{33\text{-}274}$:CO8 complex in two crystal forms. Visualization and coloration follows Figure 1B. (A) CHIP13$^{33\text{-}274}$:CO8, Crystal form 1 (PDB: 9Q84). (B) CHIP13$^{33\text{-}274}$:CO8, Crystal form 2 (PDB: 9Q83) (C) Surface representation of CHIP13:CO8 binding mode. CO8 is displayed in ball-and-stick representation. The closed groove of LysM2 and the open groove of LysM3 are indicated. (D) Detailed representation of the CO8 binding mode. LysM1 was omitted from the visualization for clarity. Residues participating in CO8 binding are indicated. F167 blocking the distal LysM2 groove is highlighted in cyan. (E) Superposition of CHIP13 and CHIP13:CO8 crystal structures. Root mean square deviation between Cα atoms, 0.263 Å. (F) Modelling of the CHIP13 F232A Y233A binding site mutant. The alanine substitutions (in bold) spatially allow for chitin binding but abolish the CH- π interactions.

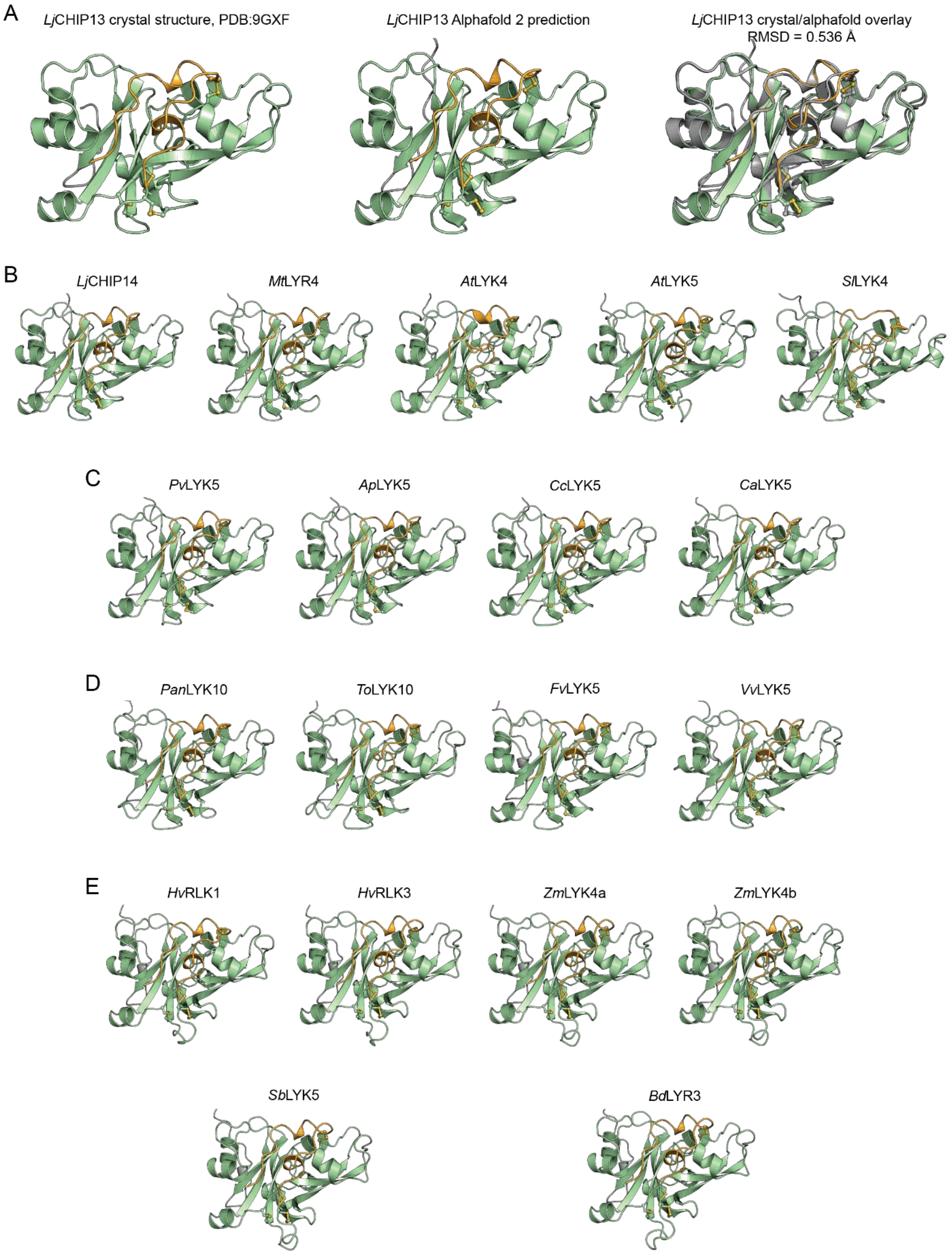
A
LjCHIP13 crystal structure, PDB:9GXF
LjCHIP13 Alphafold 2 prediction
LjCHIP13 crystal/alphafold overlay
RMSD = 0.536 Å
B
LjCHIP14
MtLYR4
AtLYK4
AtLYK5
SlLYK4
C
PvLYK5
ApLYK5
CcLYK5
CaLYK5
D
PanLYK10
ToLYK10
FvLYK5
VvLYK5
E
HvRLK1
HvRLK3
ZmLYK4a
ZmLYK4b
SbLYK5
BdLYR3

**Fig. S3. The bridge domain can be structurally predicted in LysM-RKs from other species.** A-E: The bridge domain of CHIP13 and the additional disulphide bridge can be predicted in AlphaFold2. The bridge domain is colored orange; disulphide bridges are drawn in stick representation. (A) AlphaFold2 model of CHIP13 compared to experimental crystal structure. Left panel: Experimental structure of CHIP13 without ligand, PDB 9GXF. Middle panel: AlphaFold2 model of *Lotus japonicus* CHIP13 ectodomain. Uniprot Entry D3KU00_LOTJA. Right panel: Superposition of CHIP13 experimental structure and AlphaFold2 prediction showing high structural similarity. Alpha carbon r.m.s.d = 0.536 Å. (B) AlphaFold2 models of established chitin receptors of the CHIP13-class. *Lotus japonicus* CHIP14: D3KU01_LOTJA ; *Medicago truncatula* LYR4: G7K7K0_MEDTR ; *Arabidopsis thaliana* LYK4: O64825/LYK4_ARATH ; *Arabidopsis thaliana* LYK5: O22808/ LYK5_ARATH; *Solanum lycopersicum* LYK4: XP_004232089 (C) CHIP13 homologues from other legumes: *Phaseolus vulgaris* LYK5: V7BAY6_PHAVU; *Abrus precatorius* LYK5: A0A8B8JHM2_ABRPR; *Cajanus cajan* LYK5 A0A151TVG1_CAJCA; Cicer arietinum LYK5: A0A1S2XLD6_CICAR (D) CHIP13 homologues from other dicots: *Parasponia andersonii* LYK10: A0A2P5AHA4_PARAD; *Trema orientale* LYK5: A0A221I0Y2_TREOI; *Fragaria vesca* LYK5: XP_004301882 ;*Vitis vinifera* LYK5: F6I6Y0_VITVI (E) CHIP13 homologues from monocots: *Hordeum vulgare* RLK1: F2E2C9_HORVV ; *Hordeum vulgare* RLK3: F2E7K4_HORVV ; *Zea mays* LYK4a: B6SWT1 ; *Zea mays* LYK4b: A0A804PG17; *Sorghum bicolor* LYK5: C5XXD8_SORBI; *Brachypodium distachyon* LYK5: I1HY83_BRADI.

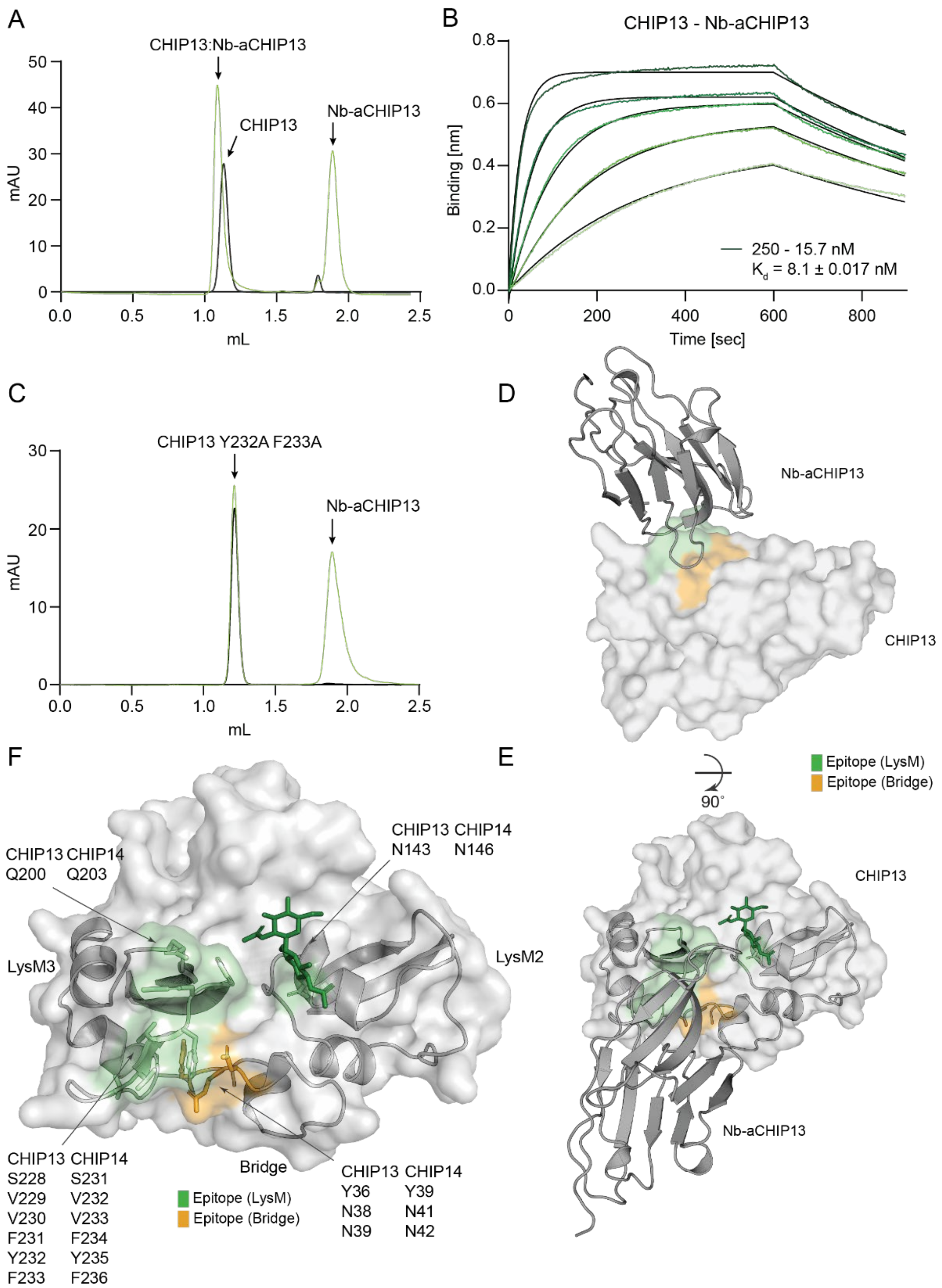

A
CHIP13:Nb-aCHIP13
CHIP13
Nb-aCHIP13
mAU
mL
B
CHIP13 - Nb-aCHIP13
Binding [nm]
Time [sec]
250 - 15.7 nM
K_d = 8.1 ± 0.017 nM
C
CHIP13 Y232A F233A
Nb-aCHIP13
mAU
mL
D
Nb-aCHIP13
CHIP13
F
CHIP13 CHIP14
Q200 Q203
CHIP13 CHIP14
N143 N146
LysM3
LysM2
CHIP13 CHIP14
S228 S231
V229 V232
V230 V233
F231 F234
Y232 Y235
F233 F236
Bridge
Epitope (LysM)
Epitope (Bridge)
CHIP13 CHIP14
Y36 Y39
N38 N41
N39 N42
E
Epitope (LysM)
Epitope (Bridge)
90°
CHIP13
Nb-aCHIP13

**Fig. S4. Nb-aCHIP13 binds CHIP13 with high affinity and the epitope is conserved in CHIP14.**

(A) $A_{280}$ chromatogram of SEC analysis of stable Nb-aCHIP13:CHIP13 complex on an analytical Superdex 75 increase, 3.2/300 column. The CHIP13:Nb-aCHIP13 complex elutes earlier than CHIP13 alone. (B) BLI sensorgram of Nb-aCHIP13:CHIP13 complex for affinity and kinetics determination. CHIP13 binds Nb-aCHIP13 with high affinity (Kd = 8.1 ± 0.017 nM). (C) $A_{280}$ chromatogram of SEC analysis testing Nb-aCHIP13 binding to CHIP13 Y232A F233A on an analytical Superdex 75 increase, 3.2/300 column. (D-F) Mapping of the Nb-aCHIP13 epitope and its conservation in CHIP14. (D) Crystal structure of the CHIP13:Nb-aCHIP13 complex, same view as Figure 2F. Nb-aCHIP13 is displayed as cartoon, CHIP13 in surface representation. The surface area contacted by the nanobody is displayed in color (orange: bridge domain; green: LysM core). (E) Same representation as in panel D, turned 90°, showing the full epitope. (F) Detailed visualization of CHIP13 epitope; residues involved in nanobody binding are highlighted in stick representation. The glycan on N143, which is directly bound by the nanobody, is displayed in dark green. Core LysM2, LysM3 and the bridge domain are shown in cartoon representation. Labels indicate amino acids on CHIP13 and CHIP14, respectively. Nb-aCHIP13 was omitted from the visualization.

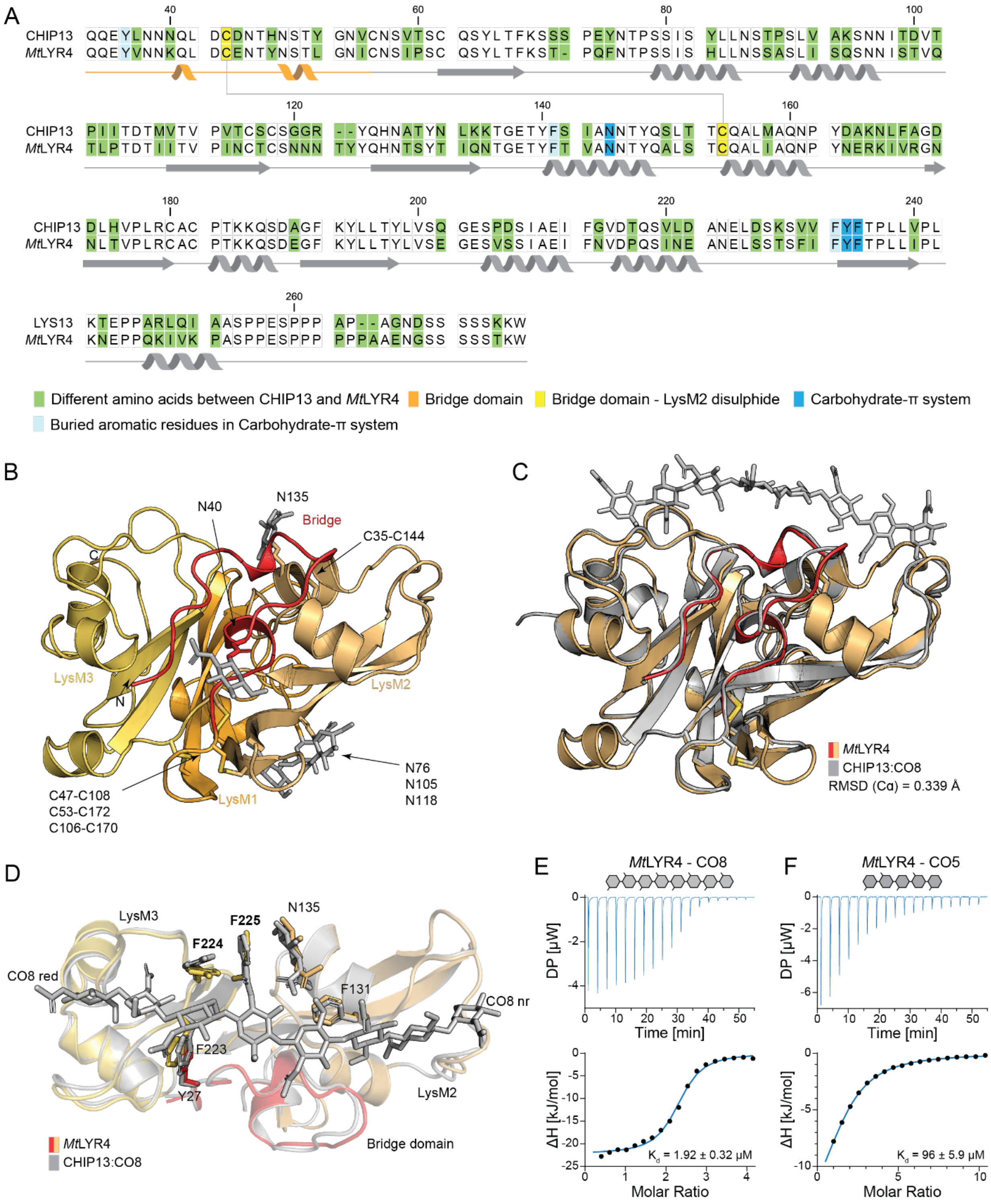
A
40 60 80 100
CHIP13 QQEYLNNNQL DCDNTHNSTY GNVCNSVTSC QSYLTFKSSS PEYNTPSSIS YLLNSTPSLV AKSNNITDVT
MtLYR4 QQEYVNNKQL DCENTYNSTL GNICNSIPSC QSYLTFKST- PQFNTPSSIS HLLNSSASLI SQSNNISTVQ
120 140 160
CHIP13 PIITDTMVTV PVTCSCSGGR --YQHNATYN LKKTGETYFS IANNTYQSLT TCQALMAQNP YDAKNLFAGD
MtLYR4 TLPTDTIITV PINCTCSNNN TYYQHNTSYT IQNTGETYFT VANNTYQALS TCQALIAQNP YNERKIVRGN
180 200 220 240
CHIP13 DLHVPLRCAC PTKKQSDAGF KYLLTYLVSQ GESPDSIAEI FGVDTQSVLD ANELDSKSVV FYFTPLLVPL
MtLYR4 NLTVPLRCAC PTKKQSDEGF KYLLTYLVSE GESVSSIAEI FNVDPQSINE ANELSSTSFI FYFTPLLIPL
260
LYS13 KTEPPARLQI AASPPESPPP AP--AGNDSS SSSKKW
MtLYR4 KNEPPQKIVK PASPPESPPP PPPAAENGSS SSSTKW
Different amino acids between CHIP13 and MtLYR4
Bridge domain
Bridge domain - LysM2 disulphide
Carbohydrate-π system
Buried aromatic residues in Carbohydrate-π system
B
N40
N135
Bridge
C35-C144
C
LysM3
N
LysM2
N76
N105
N118
C47-C108
C53-C172
C106-C170
LysM1
C
MtLYR4
CHIP13:CO8
RMSD (Cα) = 0.339 Å
D
LysM3
F225
N135
F224
CO8 red
F131
CO8 nr
F223
Y27
LysM2
MtLYR4
CHIP13:CO8
Bridge domain
E
MtLYR4 - CO8
DP [μW]
Time [min]
ΔH [kJ/mol]
Kd = 1.92 ± 0.32 μM
Molar Ratio
F
MtLYR4 - CO5
DP [μW]
Time [min]
ΔH [kJ/mol]
Kd = 96 ± 5.9 μM
Molar Ratio

**Fig. S5. *Mt*LYR4 is conserved in structure and function to CHIP13/14.**
(A) Sequence alignment of CHIP13 and *Mt*LYR4 ectodomains with highlighted bridge domain (orange), additional disulfide bridge (yellow) and aromatic motif. Dark blue: exposed aromatic residues and N-glycan; light blue: buried aromatic residues associated with carbohydrate-π network. Green: Amino acid differences between CHIP13 and LYR4. The predicted signal peptide was omitted from the sequence alignment. (B) Cartoon representation of the crystal structure of *Mt*LYR4 with the N-terminal domain and the additional disulfide bridge C35-C144. Glycans are depicted in light gray stick representation. (C) Superposition of CHIP13:CO8 and LYR4 crystal structures. Alpha carbon RMSD = 0.339 Å between the two structures. (D) Superposition of CHIP13:CO8 and LYR4 crystal structures, zoom into the ligand binding site. Residues participating in carbohydrate-π interactions and CO8 from CHIP13 are highlighted in stick representation. F224 and F225 are highlighted in bold. LysM1 was omitted for clarity. (E) ITC isotherms for titration of CO8 (left) and CO5 (right) into LYR4.

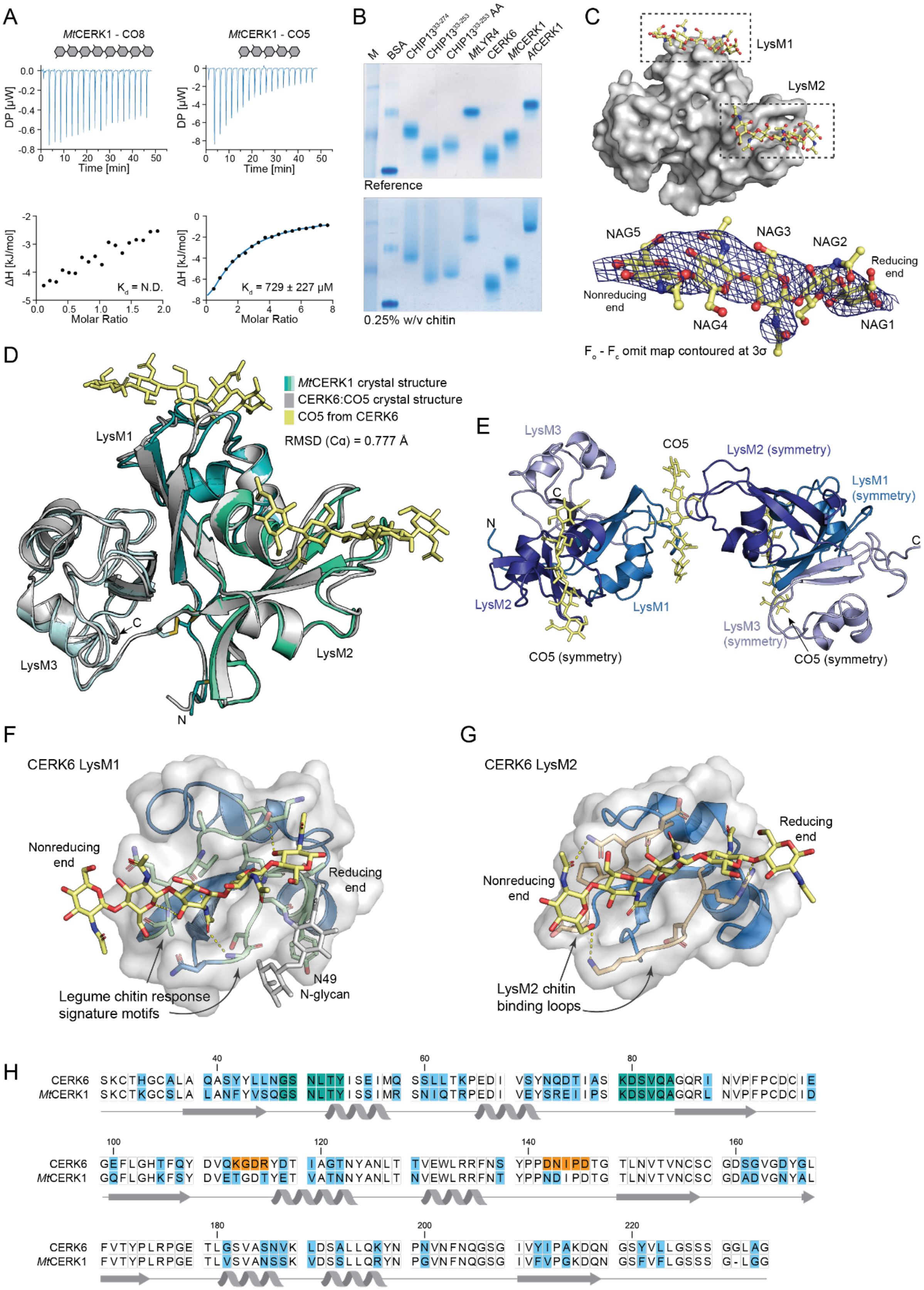

A
MtCERK1 - CO8
MtCERK1 - CO5
DP [μW]
Time [min]
ΔH [kJ/mol]
Molar Ratio
K_d = N.D.
K_d = 729 ± 227 μM
B
M
BSA
MtLYR4
CERK6
MtCERK1
AtCERK1
Reference
0.25% w/v chitin
C
LysM1
LysM2
NAG5
NAG3
NAG2
Reducing end
Nonreducing end
NAG4
NAG1
F_o - F_c omit map contoured at 3σ
D
MtCERK1 crystal structure
CERK6:CO5 crystal structure
CO5 from CERK6
RMSD (Cα) = 0.777 Å
LysM1
LysM2
LysM3
C
N
E
LysM3
CO5
LysM2 (symmetry)
LysM1 (symmetry)
N
C
LysM2
LysM1
LysM3 (symmetry)
CO5 (symmetry)
CO5 (symmetry)
F
CERK6 LysM1
Nonreducing end
Reducing end
N49 N-glycan
Legume chitin response signature motifs
G
CERK6 LysM2
Reducing end
Nonreducing end
LysM2 chitin binding loops
H
CERK6 SKCTHGCALA QASYYLLNGS NLTYISEIMQ SSLLTKPEDI VSYNQDTIAS KDSVQAGQRI NVPFPCDCIE
MtCERK1 SKCTKGCSLA LANFYVSQGS NLTYISSIMR SNIQTRPEDI VEYSREIIPS KDSVQAGQRL NVPFPCDCID
CERK6 GEFLGHTFQY DVQKGDRYDT IAGTNYANLT TVEWLRRFNS YPPDNIPDTG TLNVTVNCSC GDSGVGDYGL
MtCERK1 GQFLGHKFSY DVETGDTYET VATNNYANLT NVEWLRRFNT YPPNDIPDTG TLNVTVNCSC GDADVGNYAL
CERK6 FVTYPLRPGE TLGSVASNVK LDSALLQKYN PNVNFNQGSG IVYIPAKDQN GSYVLLGSSS GGLAG
MtCERK1 FVTYPLRPGE TLVSVANSSK VDSSLLQRYN PGVNFNQGSG IVFVPGKDQN GSFVFLGSSS G-LGG

**Fig. S6. CERK6 and *Mt*CERK1 are weak chitin binders.**
(A) ITC binding isotherm of CO8 (left) and CO5 (right) titration into *Mt*CERK1. Measurement of CO8 binding to *Mt*CERK1 is limited by CO8 solubility and $K_d$ cannot be accurately determined (> 5 mM). (B) Affinity PAGE assay showing retention of ectodomains in chitin gel. Left: Control Native PAGE; Right: Affinity PAGE containing 0.25% (w/v) shrimp shell chitin. (C) $F_o - F_c$ omit map of CO5 electron density contoured at 3σ. Resolution is d = 3 Å. Top panel shows CO5 as ball-and-sticks bound by CERK6 in surface representation. Dashed box indicates symmetry-related ligands and binding sites. (D) Superposition of CERK6:CO5 (gray cartoon) and MtCERK1 crystal structures. CO5 from the CERK6:CO5 structure is displayed in yellow stick representation. Alpha carbon RMSD = 0.777 Å between the two crystal structures. (E) The crystallographic sandwich-type chitin binding in the CERK6 crystal packing. CERK6 binds CO5 in both LysM1 and LysM2, each shared with a symmetry-related CERK6. (F) Detailed view of CERK1:CO5 binding in LysM1. CO5 is displayed in yellow stick representation and polar contacts indicated. Conserved legume signature motifs are highlighted in green. The glycan on N46 is drawn in gray sticks. (G) Detailed view of CERK1:CO5 binding in LysM2. Chitin binding loops and polar contacts are indicated. (H) Sequence conservation of CERK6 and MtCERK1 ectodomains. Diverging residues are highlighted in light blue. Conserved legume LysM1 signature motifs are highlighted in teal, LysM2 chitin binding residues in orange. Indicated secondary structure elements are based on the crystal structures. The signal peptide was omitted from the alignment.

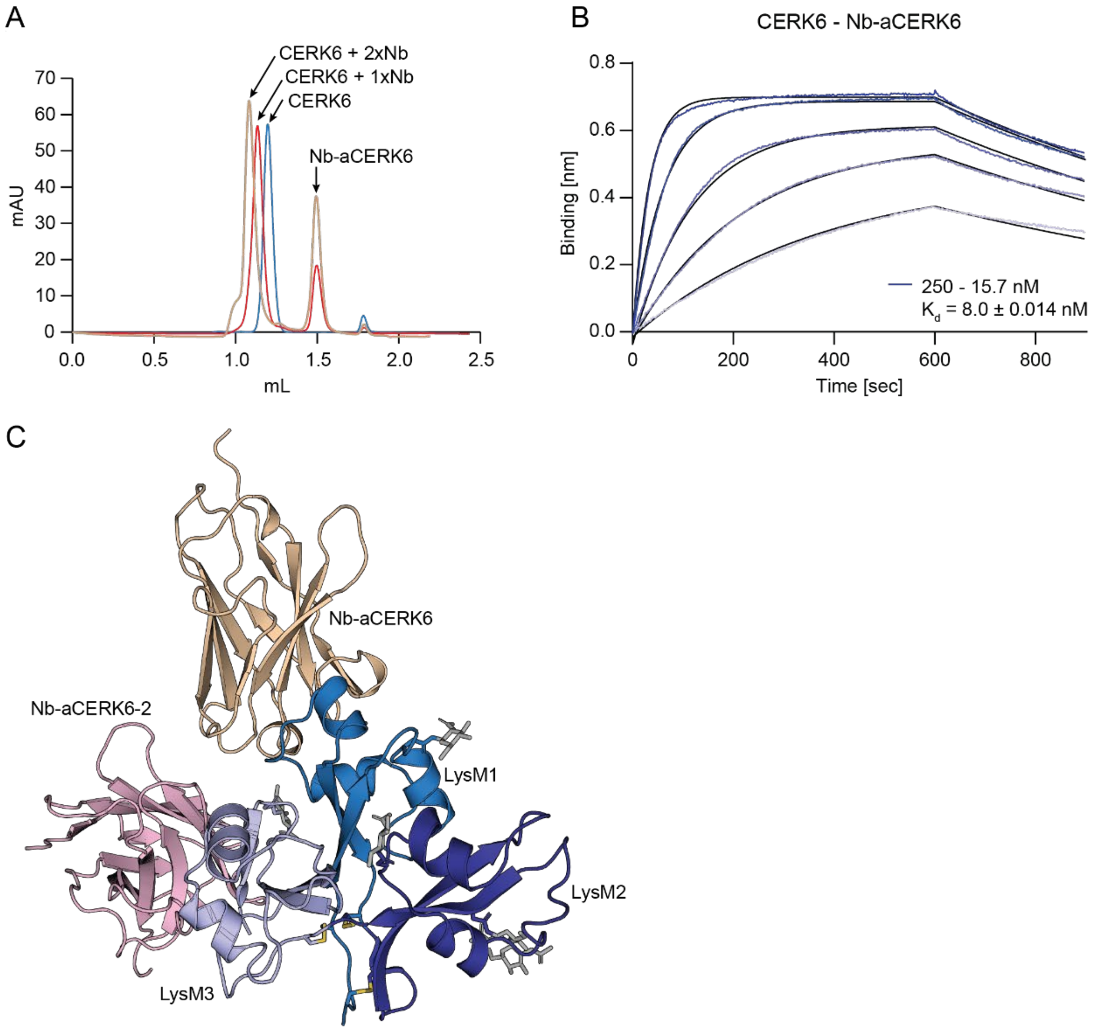
A
CERK6 + 2xNb
CERK6 + 1xNb
CERK6
Nb-aCERK6
mAU
mL
B
CERK6 - Nb-aCERK6
Binding [nm]
Time [sec]
250 - 15.7 nM
Kd = 8.0 ± 0.014 nM
C
Nb-aCERK6
Nb-aCERK6-2
LysM1
LysM2
LysM3

**Fig. S7. Specific nanobodies against CERK6 ectodomain.**
(A) $A_{280}$ SEC chromatogram of stable Nb-aCERK6:CERK6 (red) and Nb-aCERK6:Nb-aCERK6-2:CERK6 (beige) complex on an analytical Superdex 75 Increase, 3.2/300 column in phosphate-buffered saline, pH 7.4. (B) BLI sensorgram of CERK6:Nb-aCERK6 binding. (D) Total view of the crystal structure of CERK6 in complex with two nanobodies, Nb-aCERK6 and Nb-aCERK6-2.

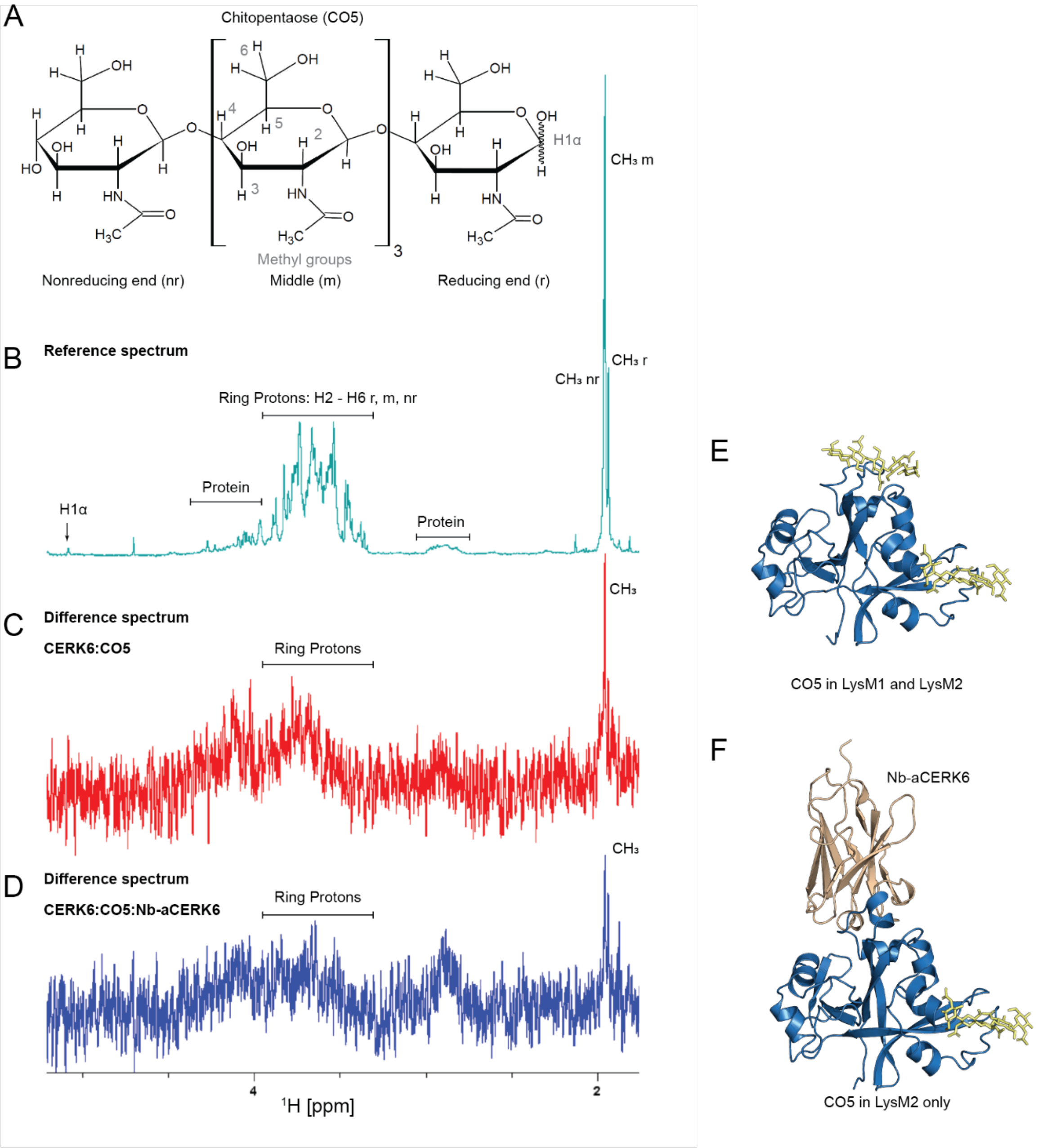

A
Chitopentaose (CO5)
Methyl groups
Nonreducing end (nr)
Middle (m)
Reducing end (r)
H1α
CH₃ m
B
Reference spectrum
CH₃ r
CH₃ nr
Ring Protons: H2 - H6 r, m, nr
Protein
H1α
Protein
C
Difference spectrum
CERK6:CO5
Ring Protons
CH₃
D
Difference spectrum
CERK6:CO5:Nb-aCERK6
Ring Protons
CH₃
4
2
¹H [ppm]
E
CO5 in LysM1 and LysM2
F
Nb-aCERK6
CO5 in LysM2 only

**Fig. S8. CERK6 can accommodate chitin in LysM1 and LysM2.**

STD-NMR analysis of CERK6:CO5 and CERK6:Nb-aCERK6:CO5 complexes. (A) Structural formula of CO5 with relevant protons indicated. (B) Reference spectrum (C) Difference spectrum of CERK6:CO5 (20 µM:1 mM); (D) CERK6:Nb-aCERK6:CO5 (20 µM: 1 mM). Both difference spectra show binding to the methyl groups of CO5 as well as H2-H6 in both nonreducing end, middle and reducing end carbohydrate rings. (E) CERK6 with chitin in LysM1 and LysM2, binding mode seen in spectrum shown in (C), (F) CERK6 with chitin in LysM2 and Nb-aCERK6 in LysM1, binding mode of spectrum shown in (D).

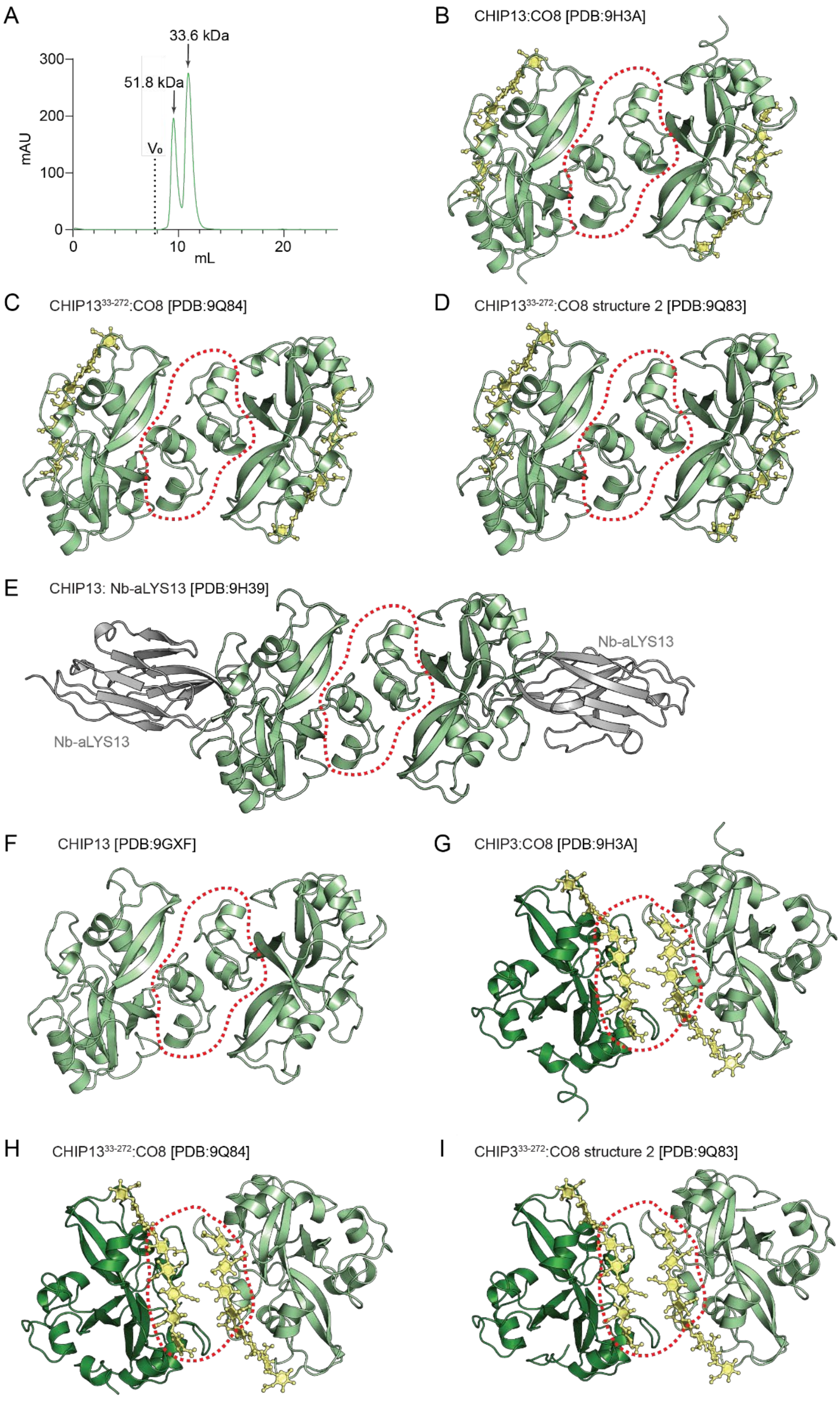

A
33.6 kDa
51.8 kDa
V₀
mAU
300
200
100
0
0
10
20
mL
B
CHIP13:CO8 [PDB:9H3A]
C
CHIP13^33-272:CO8 [PDB:9Q84]
D
CHIP13^33-272:CO8 structure 2 [PDB:9Q83]
E
CHIP13: Nb-aLYS13 [PDB:9H39]
Nb-aLYS13
Nb-aLYS13
F
CHIP13 [PDB:9GXF]
G
CHIP3:CO8 [PDB:9H3A]
H
CHIP13^33-272:CO8 [PDB:9Q84]
I
CHIP3^33-272:CO8 structure 2 [PDB:9Q83]

**Fig. S9. Different conserved interfaces in CHIP13 crystals.**
(A) CHIP13 elutes in dimer (~51 kDa) and monomer (~33 kDa) populations from SEC at high concentrations. SEC profile of CHIP13 without ligand on a Superdex 75 increase 10/300 GL column in phosphate buffered saline, pH 7.4. Arrows denote calibrated molecular mass, the dashed line the void volume $V_o$ as determined by blue dextran calibration. (B-F) Ligand-independent dimer formation mediated by a LysM1-LysM1 interaction present in all our CHIP13 crystal structures. The dimer interface is highlighted with a dashed line. CHIP13 is displayed in pale green, the CO8 ligand, if present, in yellow stick representation (B) Crystal structure of CHIP13:CO8 (9H3A). The ligand-independent dimer is comprised of two symmetry-related molecules (C) CHIP13$^{33\text{-}274}$, crystal form 1 (9Q84). Two symmetry-related molecules form the dimer. (D) LysM1-LysM1 dimer in CHIP13$^{33\text{-}274}$:CO8, crystal form 2 (9Q83). Two symmetry-related molecules form the interaction. (E) CHIP13:Nb-aCHIP13 (9H39). The dimer constitutes the asymmetric unit. (F) Crystal structure of CHIP13 without ligand (9GXF). The dimer constitutes the asymmetric unit. (G-I) Ligand-induced dimer formation mediated by CO8 in all our CHIP13 crystal structures with CO8. (G) Crystal structure of CHIP13:CO8 (9H3A). The ligand-induced dimer is comprised of two symmetry-related molecules. (H) CHIP13$^{33\text{-}274}$, crystal form 1 (9Q84). Two symmetry-related molecules constitute the ligand-induced dimer (I) CHIP13$^{33\text{-}274}$, crystal form 2 (9Q83). The ligand-induced dimer constitutes the asymmetric unit.

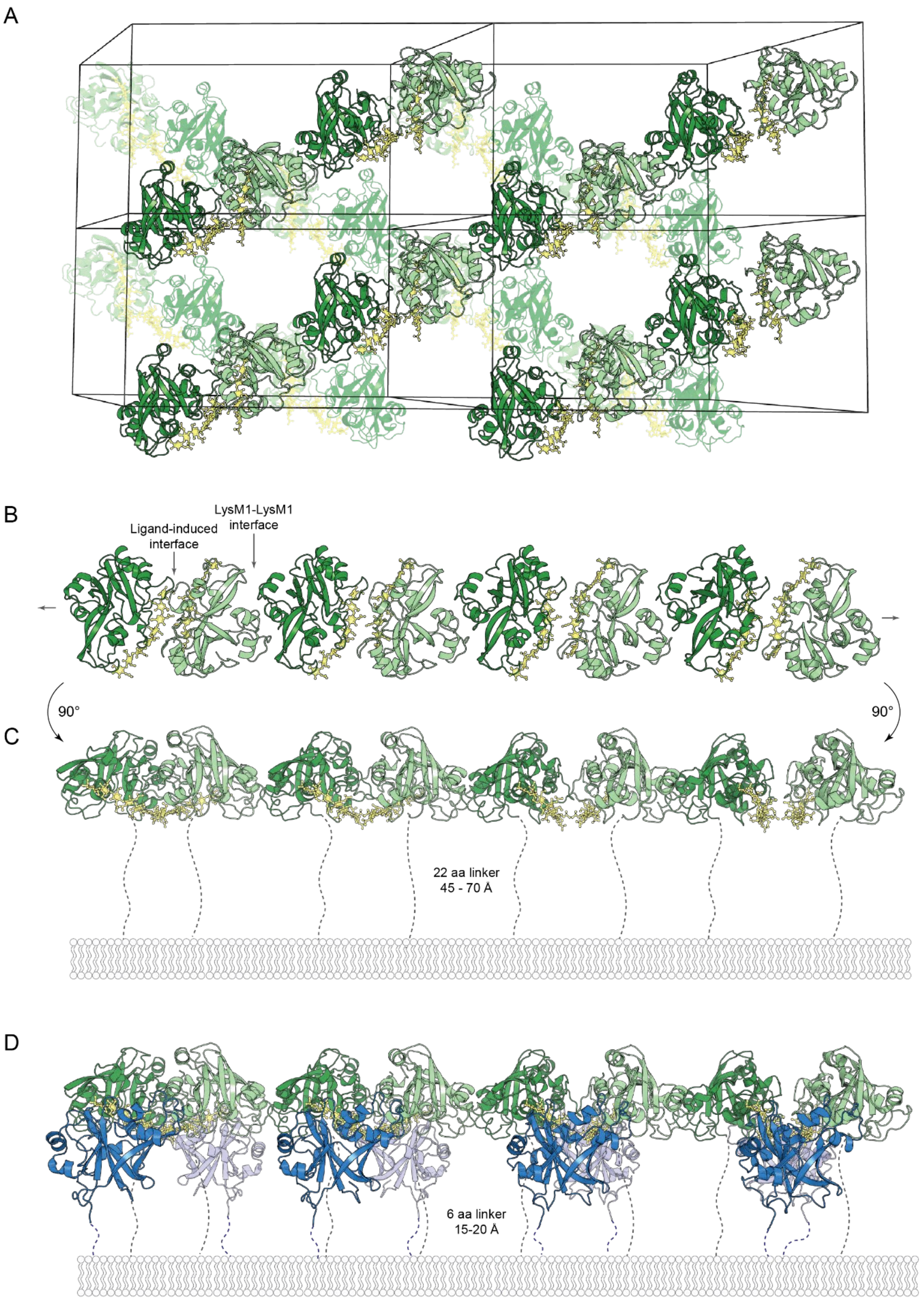
A
B
Ligand-induced interface
LysM1-LysM1 interface
90°
90°
C
22 aa linker
45 - 70 Å
D
6 aa linker
15-20 Å

**Fig. S10. Crystal packing analysis of the CHIP13:CO8 complex and docking of the active chitin perception complex.**

(A) Crystal packing of the CHIP13$^{33\text{-}274}$:CO8 crystal (PDB: 9Q83). Four unit cells and their content are illustrated. CHIP13 monomers are alternating shown in dark green and light green cartoon representation, CO8 in golden ball-and-stick representation The asymmetric unit consists of two CHIP13$^{33\text{-}274}$ forming the ligand-induced dimer. CHIP13$^{33\text{-}274}$ is arranged in loosely packed rows alternating between ligand-independent (LysM1-LysM1) and ligand-induced crystal contacts. For clarity, the second layer is displayed semi-transparently. (B) Detailed view of a single crystal packing row (4 asymmetric units) consisting of CHIP13$^{33\text{-}274}$ arranged in an alternating ligand-independent/ligand-induced scaffold. (C) 90 ° turned side view of B. The C-termini of CHIP13$^{33\text{-}274}$ , their possible orientation and spacing towards the plasma membrane drawn in dashed lines. (D) *In silico* model of the active chitin perception heterocomplex after recruitment of CERK6. CO5 bound in LysM1 of CO5 was superimposed with the fixed nonreducing end of the CO8 bound by CHIP13 LysM2. Possible orientation, length and spacing of the C-termini of all receptor ectodomains are indicated with dashed lines.

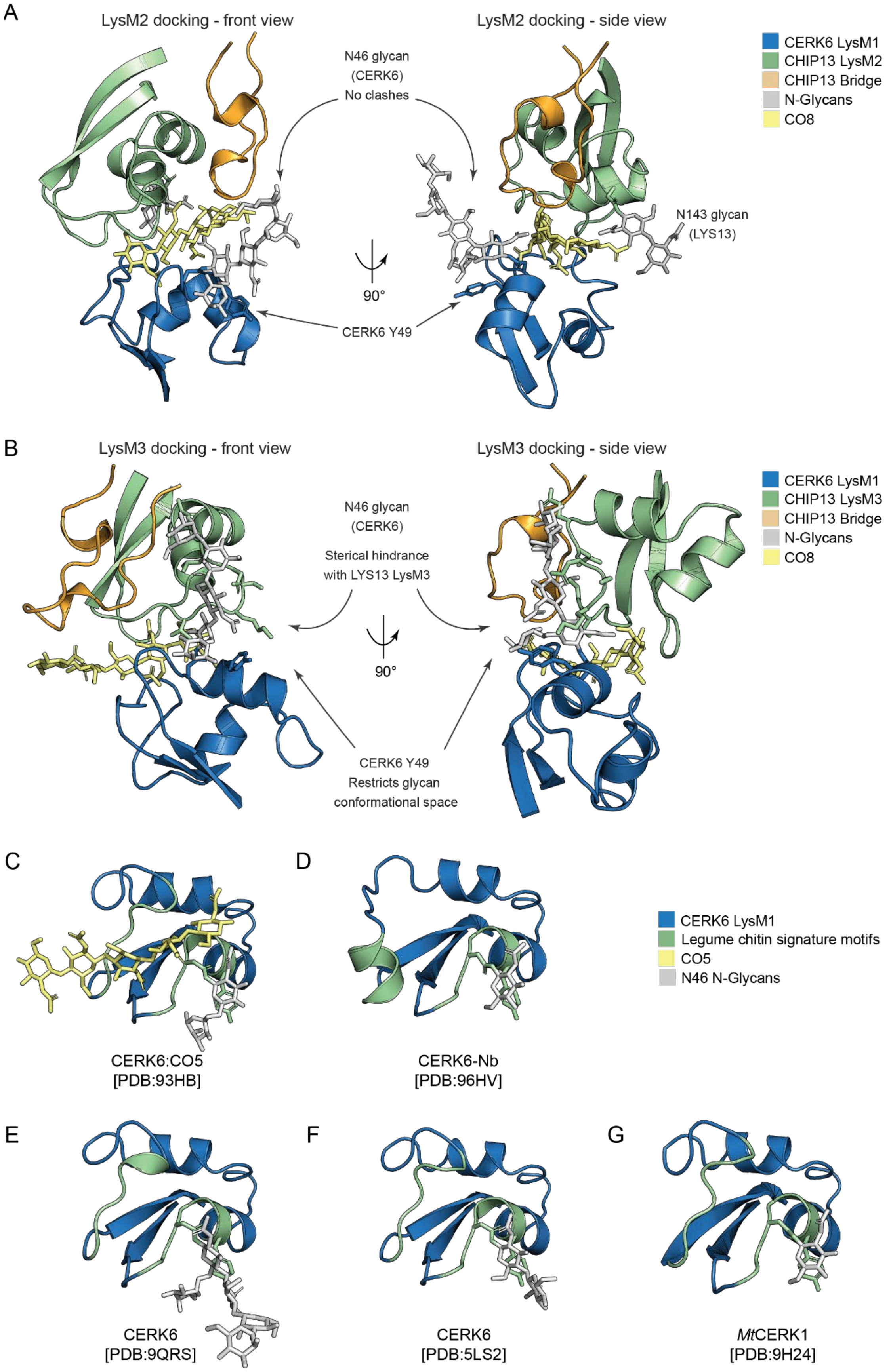
A
LysM2 docking - front view
LysM2 docking - side view
CERK6 LysM1
CHIP13 LysM2
CHIP13 Bridge
N-Glycans
CO8
N46 glycan
(CERK6)
No clashes
N143 glycan
(LYS13)
90°
CERK6 Y49
B
LysM3 docking - front view
LysM3 docking - side view
CERK6 LysM1
CHIP13 LysM3
CHIP13 Bridge
N-Glycans
CO8
N46 glycan
(CERK6)
Sterical hindrance
with LYS13 LysM3
90°
CERK6 Y49
Restricts glycan
conformational space
C
D
CERK6 LysM1
Legume chitin signature motifs
CO5
N46 N-Glycans
CERK6:CO5
[PDB:93HB]
CERK6-Nb
[PDB:96HV]
E
F
G
CERK6
[PDB:9QRS]
CERK6
[PDB:5LS2]
MtCERK1
[PDB:9H24]

**Fig. S11.** ***In silico*** **docking of the CHIP13-CERK6 complex.**
(A) *In silico* model of ligand-induced heterodimer formation between CERK6 LysM1 and CHIP13 LysM2 and bridge domain. CERK6 LysM1, CHIP13 LysM2 and bridge domain are visualized as cartoons, the CERK6 N46, CHIP13 N143 N-glycans and CO8 are drawn as sticks. Remaining receptor parts are omitted from the representation. (B) *In silico* model of ligand-induced heterodimer formation between CERK6 LysM1 and CHIP13 LysM3 and bridge domain. CERK6 LysM1, CHIP13 LysM3 and bridge domain are visualized as cartoons, the CERK6 N46 N-glycans and CO8 are drawn as sticks. CERK6 Y49 (part of the conserved legume signature motif) and CHIP13 residues 223-229 are indicated in stick representation. The conserved CERK6 Y49 interacts with the glycan core and further restricts its conformational space (C-G) The N-glycan on N46 is present in all crystal structures of CERK6 and MtCERK1. Cartoon representation of CERK6 LysM1 with conserved legume signature motifs in green. The N46 glycan is drawn in gray sticks, the conserved Y49 in green sticks. (C) CERK6:CO5, PDB:93HB (this study). (D) CERK6-Nb, PDB: 96HV (this study). (E) CERK6 without ligand, PDB: 9QRS (this study). (F) CERK6 without ligand, PDB 5LS2 (*3*). (G) MtCERK1, PDB: 9H24 (this study).

**Table S1. (separate file)**

Data collection and refinement statistics for all crystal structures. Values in parentheses represent the highest resolution shell. For datasets where ellipsoidal processing was performed, corresponding spherical processing statistics are shown alongside (9Q83, 9H3B, 9H6V).

**Table S2. Amino acid sequences of recombinant protein expression constructs.**
Mutations are highlighted in red; protein modifications (e.g. purification tags) are highlighted in blue.

| |
|---|
| **CHIP13(33-253)**<br>MVSAIVLYVLLAAAAHSAFAQQEYLNNNQLDCDNTHNSTYGNVCNSVTSCQSYLT-FKSSSPEYNTPSSISYLLNSTPSLVAKSNNITDVTPIITDTMVTVPVTCSCSGGRYQHNATYNLKKTGETYFSIANNTYQSLTTCQALMAQNPYDAKNLFAGDDLHVPLRCACPTKKQSDAGFK-YLLTYLVSQGESPDSIAEIFGVDTQSVLDANELDSKSVVFYFTPLLVPLKTEPPARLQIAASHHHHHH |
| **CHIP13(33-274)**<br>MVSAIVLYVLLAAAAHSAFAQQEYLNNNQLDCDNTHNSTYGNVCNSVTSCQSYLT-FKSSSPEYNTPSSISYLLNSTPSLVAKSNNITDVTPIITDTMVTVPVTCSCSGGRYQHNATYNLKKTGETYFSIANNTYQSLTTCQALMAQNPYDAKNLFAGDDLHVPLRCACPTKKQSDAGFK-YLLTYLVSQGESPDSIAEIFGVDTQSVLDANELDSKSVVFYFTPLLVPLKTEPPARLQIAASPPESPPPAPAGNDSSSSSKKWHHHHHH |
| **LYR4 (24-242)**<br>MVSAIVLYVLLAAAAHSAFAQQEYVNNKQLDCENTYNSTLGNICNSIPSCQSYLT-FKSTPQFNTPSSISHLLNSSASLISQSNNISTVQTLPTDTIITVPINCTCSNNNTYYQHNTSYTIQNTGETYFTVANNTYQALSTCQALIAQNPYNERKIVRGNNLTVPLRCACPTKKQSDEGFK-YLLTYLVSEGESVSSIAEIFNVDPQSINEANELSSTSFIFYFTPLLIPLKNEPPQKIVKHHHHHH |
| **LYS13 (33-253) F232A F233A**<br>MVSAIVLYVLLAAAAHSAFAQQEYLNNNQLDCDNTHNSTYGNVCNSVTSCQSYLT-FKSSSPEYNTPSSISYLLNSTPSLVAKSNNITDVTPIITDTMVTVPVTCSCSGGRYQHNATYNLKKTGETYFSIANNTYQSLTTCQALMAQNPYDAKNLFAGDDLHVPLRCACPTKKQSDAGFK-YLLTYLVSQGESPDSIAEIFGVDTQSVLDANELDSKSVVFAATPLLVPLKTEPPARLQIAASHHHHHH |
| **MtCERK1 (30-232)**<br>MVSAIVLYVLLAAAAHSAFAKCTKGCSLALANFYVSQGSNLTYISSIMRSNIQTRPEDI-VEYSREIIPSKDSVQAGQRLNVPFPCDCIDGQFLGHKFSYDVETGDTYETVATNNYANLTNVEWLRRFNTYPPNDIPDTGTLNVTVNCSCGDADVGNYALFVTY-PLRPGETLVSVANSSKVDSSLLQRYNPGVNFNQGSGIVFVPGKDQNGSFVFLGSSSGLGGHHHHHH |
| **Nb-aCERK6**<br>MQRQLVESGGGLVQPGGSLRLSCAASGRTFSASTMGWFRQAPGKEREFVVCVSRNGESTY-YADSVKGRFIISRDNVKNTVYLQMNSLEPEDTAVYYCAARTRGIVCDSTDSYGYWGKGTPVTVSSLEHHHHHH |
| **Nb-aCERK6-2**<br>MQVQLVETGGGLVQAGGSLRLSCAASGRIFSRTIMAWFRQAPGKEREFVAAIRWSGG-DTYYTDSMKGRFTVSRDNVKNTLYLQIDSLKPEDTAVYYCAAHRLDDEARLLPASVYDYWGRGTQVTVSSHHHHHH |

| |
|---|
| **Nb-aLYS13**<br>MQLQLVESGGGLVQAGGSLRLSCATSGTTFRLNTMGWYRQAPGKQRELVATISRDFKTN-<br>YADSVKGRFTISRDNAKHTVDLQMNSLTPEDTAVYYCLVRDQREWYGPEYDNWGRGTQVTVSSHH<br>HHHH |

**Table S3. Amino acid sequences of receptors for *Medicago* complementation assays.**
Mutations are highlighted in red; protein modifications (e.g. affinity tags) are highlighted in blue.

| |
|---|
| **MtLYR4-His**<br>MAWQTLTTLIIITIITTFPKTKSQQEYVNNKQLDCENTYNSTLGNICNSIPSCQSYLT-<br>FKSTPQFNTPSSISHLLNSSASLISQSNNISTVQTLPTDTIITVPINCTCSNNNTYYQHNTSYTI<br>QNTGETYFTVANNTYQALSTCQALIAQNPYNERKIVRGNNLTVPLRCACPTKKQSDEGFK-<br>YLLTYLVSEGESVSSIAEIFNVDPQSINEANELSSTSFIFYFTPLLIPLKNEPPQKIVKPASPPE<br>SPPPPPPAAENGSSSSSTKWVIVGVVVGVVVLLLVGVALFFLCFRRRRQQKLQP-<br>PAVGKAFSDSNTKKVSEVTSTSQSWSLSSEGIRYAVDSLTVYKYEDLQNATNFFSEENKIKGSVY<br>RASFKGDDAAVKILKGDVSSEINILKRINHANIIRLSGFCVYKGNTYLVYEFA-<br>ENNSLDDWLHSEKNKDKNYSNSMCLSWFQRVQIAHDVADALNYLHNYANPPHVHKNLKSGNILLD<br>GKFRGKVSNFGLARVMENEGGDEGFQLTRHVIGTQGYMAPEYIENGLITPKMDVFAFGV-<br>VILELLSGREVVGSDKSNGLGDQLLASTVNQVLEGDNVREKLRGFMDPNLRDEYPLDLAFSMAEI<br>AKRCVARDLNSRPNVSEVFMILSKIQSSTLEWDPSGDLERSRSVSQVSDSRSSHHHHHH |
| **MtLYR4 Y224A F225A-His**<br>MAWQTLTTLIIITIITTFPKTKSQQEYVNNKQLDCENTYNSTLGNICNSIPSCQSYLT-<br>FKSTPQFNTPSSISHLLNSSASLISQSNNISTVQTLPTDTIITVPINCTCSNNNTYYQHNTSYTI<br>QNTGETYFTVANNTYQALSTCQALIAQNPYNERKIVRGNNLTVPLRCACPTKKQSDEGFK-<br>YLLTYLVSEGESVSSIAEIFNVDPQSINEANELSSTSFIFAATPLLIPLKNEPPQKIVKPASPPE<br>SPPPPPPAAENGSSSSSTKWVIVGVVVGVVVLLLVGVALFFLCFRRRRQQKLQP-<br>PAVGKAFSDSNTKKVSEVTSTSQSWSLSSEGIRYAVDSLTVYKYEDLQNATNFFSEENKIKGSVY<br>RASFKGDDAAVKILKGDVSSEINILKRINHANIIRLSGFCVYKGNTYLVYEFA-<br>ENNSLDDWLHSEKNKDKNYSNSMCLSWFQRVQIAHDVADALNYLHNYANPPHVHKNLKSGNILLD<br>GKFRGKVSNFGLARVMENEGGDEGFQLTRHVIGTQGYMAPEYIENGLITPKMDVFAFGV-<br>VILELLSGREVVGSDKSNGLGDQLLASTVNQVLEGDNVREKLRGFMDPNLRDEYPLDLAFSMAEI<br>AKRCVARDLNSRPNVSEVFMILSKIQSSTLEWDPSGDLERSRSVSQVSDSRSSHHHHHH |

**Table S4: Amino acid sequences of receptors for *Medicago* complementation assays.**
Mutations are highlighted in red; protein modifications (e.g. affinity tags) are highlighted in blue.

**CERK6 K350A-eGFP**

MEHPRLGFPITLLLFSFILLPSTSQSKCTHGCALAQASYYLLNGSNLTY-
ISEIMQSSLLTKPEDIVSYNQDTIASKDSVQAGQRINVPFPCDCIEGEFLGHTFQYDVQKGDRYD
TIAGTNYANLTTVEWLRRFNSYPPDNIPDTGTLNVTVNCSCGDSGVGDYGLFVTY-
PLRPGETLGSVASNVKLDSALLQKYNPNVNFNQGSGIVYIPAKDQNGSYVLLGSSSGGLAGGAIA
GIAAGVAVCLLLLAGFIYVGYFRKKRIQKEELLSQETRAIFPQDGK-
DENPRSTVNETPGPGGPAAMAGITVDKSVEFSYDELATATDNFSLANKIGQGGFGSVYYAELRGE
RAAIAKMDMQASKEFLAELKVLTRVHHLNLVRLIGYSIEGSLFLVYEFIENGNLSQHL-
RGSGRDPLPWATRVQIALDSARGLEYIHEHTVPVYIHRDIKSANILIDKNYRGKVADFGLTKLTE
VGSSSLPTGRLVGTFGYMPPEYAQYGDVSPKVDVYAFGVVLYELISAKDAIV-
KTSESITDSKGLVALFEGVLSQPDPTEDLRKLVDQRLGDNYPVDSVRKMAQLAKACTQDNPQLRP
SMRSIVVALMTLSSTTDDWDVGSFYENQNLVNLMSGRGGMVSKGEELFTGVVPILVELD-
GDVNGHKFSVSGEGEGDATYGKLTLKFICTTGKLPVPWPTLVTTLTYGVQCFSRYPDHMKQHDFF
KSAMPEGYVQERTIFFKDDGNYKTRAEVKFEGDTLVNRIELKGIDFKEDGNIL-
GHKLEYNYNSHNVYIMADKQKNGIKVNFKIRHNIEDGSVQLADHYQQNTPIGDGPVLLPDNHYLS
TQSALSKDPNEKRDHMVLLEFVTAAGITLGMDELYK

**CERK6 K350A-mCherry**

MEHPRLGFPITLLLFSFILLPSTSQSKCTHGCALAQASYYLLNGSNLTY-
ISEIMQSSLLTKPEDIVSYNQDTIASKDSVQAGQRINVPFPCDCIEGEFLGHTFQYDVQKGDRYD
TIAGTNYANLTTVEWLRRFNSYPPDNIPDTGTLNVTVNCSCGDSGVGDYGLFVTY-
PLRPGETLGSVASNVKLDSALLQKYNPNVNFNQGSGIVYIPAKDQNGSYVLLGSSSGGLAGGAIA
GIAAGVAVCLLLLAGFIYVGYFRKKRIQKEELLSQETRAIFPQDGK-
DENPRSTVNETPGPGGPAAMAGITVDKSVEFSYDELATATDNFSLANKIGQGGFGSVYYAELRGE
RAAIAKMDMQASKEFLAELKVLTRVHHLNLVRLIGYSIEGSLFLVYEFIENGNLSQHL-
RGSGRDPLPWATRVQIALDSARGLEYIHEHTVPVYIHRDIKSANILIDKNYRGKVADFGLTKLTE
VGSSSLPTGRLVGTFGYMPPEYAQYGDVSPKVDVYAFGVVLYELISAKDAIV-
KTSESITDSKGLVALFEGVLSQPDPTEDLRKLVDQRLGDNYPVDSVRKMAQLAKACTQDNPQLRP
SMRSIVVALMTLSSTTDDWDVGSFYENQNLVNLMS-
GRGGMVSKGEEDNMAIIKEFMRFKVHMEGSVNGHEFEIEGEGEGRPYEGTQTAKLKVTKGGPLPF
AWDILSPQFMYG-
SKAYVKHPADIPDYLKLSFPEGFKWERVMNFEDGGVVTVTQDSSLQDGEFIYKVKLRGTNFPSDG
PVMQKKTMGWEASSERMYPEDGALKGEIKQRLKLKDGGHYDAEVKTTYKAKKPVQLP-
GAYNVNIKLDITSHNEDYTIVEQYERAEGRHSTGGMDELYK

**LYS13-eGFP**

MLLCSSSTSITMTVLLLLLVAMSFHMISETQAQQEYLNNNQLDCDNTHNSTYG-
NVCNSVTSCQSYLTFKSSSPEYNTPSSISYLLNSTPSLVAKSNNITDVTPIITDTMVTVPVTCSC
SGGRYQHNATYNLKKTGETYFSIANNTYQSLTTCQALMAQNPYDAKNLFAGDDLHVPLRCAC-
PTKKQSDAGFKYLLTYLVSQGESPDSIAEIFGVDTQSVLDANELDSKSVVFYFTPLLVPLKTEPP
ARLQIAASPPESPPPAPAGNDSSSSSKKWVIVGVTVGVAVCLVVALLVFFLCFYN-
RRRRQPAPPPVSVKDFPDSAVKMVSETTPTTESWSLSSEGVRYAIESLTAYKFGDIQTATKFFSE
ENKIKGSVYRASFKGDDAAVKILNGDVSAEINLLKRINHANIIRLSGFCVHKGNTYLVYEFA-
ENDSLDDWLHSEKKYQNSVSLSWMQRVQIAYDVADALNYLHNYTNPVLIHKNLKSGNVLLNGKFR
AKVSNFGLARAMEDQGEDGGGFQMTRHVVGTQGYMPPEYTENGLITPKMDVYAFGVVM-
LELLSGKEATGNGDKNGLGEKMVLSETVNHVLEGDNDNVRDKLRGFMDQTLRDEYPLDLAYSMAE

IAKRCVAHDLNSRPNISEVFMTLSKVQSSTLDWDPSSEV-
ERSRSVSQISESRGGMVSKGEELFTGVVPILVELDGDVNGHKFSVSGEGEGDATYGKLTLKFICT
TGKLPVPWPTLVTTLTYGVQCFSRYPDHMKQHDFFKSAMPEGYVQERTIFFKDDGNYK-
TRAEVKFEGDTLVNRIELKGIDFKEDGNILGHKLEYNYNSHNVYIMADKQKNGIKVNFKIRHNIE
DGSVQLADHYQQNTPIGDGPVLLPDNHYLSTQSALSKDPNEKRDHMVLLEFVTAA-
GITLGMDELYK

**LYS13-mCherry**

MLLCSSSTSITMTVLLLLLVAMSFHMISETQAQQEYLNNNQLDCDNTHNSTYG-
NVCNSVTSCQSYLTFKSSSPEYNTPSSISYLLNSTPSLVAKSNNITDVTPIITDTMVTVPVTCSC
SGGRYQHNATYNLKKTGETYFSIANNTYQSLTTCQALMAQNPYDAKNLFAGDDLHVPLRCAC-
PTKKQSDAGFKYLLTYLVSQGESPDSIAEIFGVDTQSVLDANELDSKSVVFYFTPLLVPLKTEPP
ARLQIAASPPESPPPAPAGNDSSSSSKKWVIVGVTVGVAVCLVVALLVFFLCFYN-
RRRRQPAPPPVSVKDFPDSAVKMVSETTPTTESWSLSSEGVRYAIESLTAYKFGDIQTATKFFSE
ENKIKGSVYRASFKGDDAAVKILNGDVSAEINLLKRINHANIIRLSGFCVHKGNTYLVYEFA-
ENDSLDDWLHSEKKYQNSVSLSWMQRVQIAYDVADALNYLHNYTNPVLIHKNLKSGNVLLNGKFR
AKVSNFGLARAMEDQGEDGGGFQMTRHVVGTQGYMPPEYTENGLITPKMDVYAFGVVM-
LELLSGKEATGNGDKNGLGEKMVLSETVNHVLEGDNDNVRDKLRGFMDQTLRDEYPLDLAYSMAE
IAKRCVAHDLNSRPNISEVFMTLSKVQSSTLDWDPSSEV-
ERSRSVSQISESRGGMVSKGEEDNMAIIKEFMRFKVHMEGSVNGHEFEIEGEGEGRPYEGTQTAK
LKVTKGGPLPFAWDILSPQFMYG-
SKAYVKHPADIPDYLKLSFPEGFKWERVMNFEDGGVVTVTQDSSLQDGEFIYKVKLRGTNFPSDG
PVMQKKTMGWEASSERMYPEDGALKGEIKQRLKLKDGGHYDAEVKTTYKAKKPVQLP-
GAYNVNIKLDITSHNEDYTIVEQYERAEGRHSTGGMDELYK

**LYS13 Y232A F233A-eGFP**

MLLCSSSTSITMTVLLLLLVAMSFHMISETQAQQEYLNNNQLDCDNTHNSTYG-
NVCNSVTSCQSYLTFKSSSPEYNTPSSISYLLNSTPSLVAKSNNITDVTPIITDTMVTVPVTCSC
SGGRYQHNATYNLKKTGETYFSIANNTYQSLTTCQALMAQNPYDAKNLFAGDDLHVPLRCAC-
PTKKQSDAGFKYLLTYLVSQGESPDSIAEIFGVDTQSVLDANELDSKSVVFAATPLLVPLKTEPP
ARLQIAASPPESPPPAPAGNDSSSSSKKWVIVGVTVGVAVCLVVALLVFFLCFYN-
RRRRQPAPPPVSVKDFPDSAVKMVSETTPTTESWSLSSEGVRYAIESLTAYKFGDIQTATKFFSE
ENKIKGSVYRASFKGDDAAVKILNGDVSAEINLLKRINHANIIRLSGFCVHKGNTYLVYEFA-
ENDSLDDWLHSEKKYQNSVSLSWMQRVQIAYDVADALNYLHNYTNPVLIHKNLKSGNVLLNGKFR
AKVSNFGLARAMEDQGEDGGGFQMTRHVVGTQGYMPPEYTENGLITPKMDVYAFGVVM-
LELLSGKEATGNGDKNGLGEKMVLSETVNHVLEGDNDNVRDKLRGFMDQTLRDEYPLDLAYSMAE
IAKRCVAHDLNSRPNISEVFMTLSKVQSSTLDWDPSSEV-
ERSRSVSQISESRGGMVSKGEELFTGVVPILVELDGDVNGHKFSVSGEGEGDATYGKLTLKFICT
TGKLPVPWPTLVTTLTYGVQCFSRYPDHMKQHDFFKSAMPEGYVQERTIFFKDDGNYK-
TRAEVKFEGDTLVNRIELKGIDFKEDGNILGHKLEYNYNSHNVYIMADKQKNGIKVNFKIRHNIE
DGSVQLADHYQQNTPIGDGPVLLPDNHYLSTQSALSKDPNEKRDHMVLLEFVTAA-
GITLGMDELYK

**LYS13 Y232A F233A-mCherry**

MLLCSSSTSITMTVLLLLLVAMSFHMISETQAQQEYLNNNQLDCDNTHNSTYG-
NVCNSVTSCQSYLTFKSSSPEYNTPSSISYLLNSTPSLVAKSNNITDVTPIITDTMVTVPVTCSC
SGGRYQHNATYNLKKTGETYFSIANNTYQSLTTCQALMAQNPYDAKNLFAGDDLHVPLRCAC-
PTKKQSDAGFKYLLTYLVSQGESPDSIAEIFGVDTQSVLDANELDSKSVVFAATPLLVPLKTEPP
ARLQIAASPPESPPPAPAGNDSSSSSKKWVIVGVTVGVAVCLVVALLVFFLCFYN-
RRRRQPAPPPVSVKDFPDSAVKMVSETTPTTESWSLSSEGVRYAIESLTAYKFGDIQTATKFFSE
ENKIKGSVYRASFKGDDAAVKILNGDVSAEINLLKRINHANIIRLSGFCVHKGNTYLVYEFA-
ENDSLDDWLHSEKKYQNSVSLSWMQRVQIAYDVADALNYLHNYTNPVLIHKNLKSGNVLLNGKFR
AKVSNFGLARAMEDQGEDGGGFQMTRHVVGTQGYMPPEYTENGLITPKMDVYAFGVVM-
LELLSGKEATGNGDKNGLGEKMVLSETVNHVLEGDNDNVRDKLRGFMDQTLRDEYPLDLAYSMAE
IAKRCVAHDLNSRPNISEVFMTLSKVQSSTLDWDPSSEV-
ERSRSVSQISESRGGMVSKGEEDNMAIIKEFMRFKVHMEGSVNGHEFEIEGEGEGRPYEGTQTAK
LKVTKGGPLPFAWDILSPQFMYG-
SKAYVKHPADIPDYLKLSFPEGFKWERVMNFEDGGVVTVTQDSSLQDGEFIYKVKLRGTNFPSDG
PVMQKKTMGWEASSERMYPEDGALKGEIKQRLKLKDGGHYDAEVKTTYKAKKPVQLP-
GAYNVNIKLDITSHNEDYTIVEQYERAEGRHSTGGMDELYK
